# QARS spermylation directs glutamine allocation to selective translation in cancer

**DOI:** 10.64898/2026.11.20.767968

**Authors:** Li-Peng Hu, Jun Li, Xu Wang, Yan-Miao Huo, Feng Han, Hao Zhang, Xiaotong Zhang, Mu Du, Guang-Hong Su, Shan Zhang, Xing-Yuan Hong, Yan Yang, Yan-Li Zhang, Xiao-Mei Yang, Xue-Li Zhang, Shu-Heng Jiang, Qing Li, Dong-Xue Li, Zhi-Gang Zhang

**Affiliations:** State Key Laboratory of Systems Medicine for Cancer, Shanghai Cancer Institute, Ren Ji Hospital, School of Medicine, Shanghai Jiao Tong University, Shanghai, China; Department of Radiation Oncology, Cancer Institute of Jiangsu University, Affiliated Hospital of Jiangsu University, Zhenjiang, China; Department of Biliary-Pancreatic Surgery, Ren Ji Hospital, School of Medicine, Shanghai Jiao Tong University, Shanghai, China; Department of Oncology, Lianyungang Oriental Hospital, Lianyungang, China; Department of Gastrointestinal Surgery, Ren Ji Hospital, School of Medicine, Shanghai Jiao Tong University, Shanghai, China

**Keywords:** Spermylation, Aminoacyl-tRNA synthetase, Metabolic coordination, Metabolite–macromolecule interactions, Metabolite bioactivity

## Abstract

Glutamine supports protein synthesis, nucleotide biosynthesis, and mitochondrial metabolism, yet how cancer cells prioritize its use under nutrient stress remains unclear. Here, we identify spermine-dependent modification of glutaminyl-tRNA synthetase (QARS) as a mechanism that directs glutamine utilization toward translation in pancreatic ductal adenocarcinoma (PDAC). Spermine accumulates during tumor progression and promotes QARS spermylation at Q216, enhancing aminoacylation and Gln-tRNA charging without altering glutamine uptake. Ribosome profiling and synonymous recoding reveal that QARS spermylation preferentially sustains translation of glutamine codon rich metabolic transcripts, including UPP1 and UCK2, promoting uridine utilization. Genetic or peptide-mediated inhibition of Q216 spermylation redistributes glutamine-derived carbon toward competing metabolic pathways, suppresses tumor growth in organoid, xenograft, and autochthonous PDAC models, and improves survival. In human PDAC, QARS Q216 spermylation increases with disease progression and predicts poor outcome. These findings reveal metabolite-dependent regulation of aminoacyl-tRNA synthetase as an active mechanism controlling nutrient allocation during metabolic stress.

**In Brief:** Spermine-dependent QARS Q216 spermylation enhances glutaminyl-tRNA charging, favors glutamine allocation to translation, and sustains a metabolic translation program that supports pancreatic cancer g*ro*wth.

## Introduction

Rapid tumor growth requires metabolic adaptations that allow cancer cells to thrive despite limited nutrient availability^1,2^. Glutamine is central to this process, supplying carbon and nitrogen for the tricarboxylic acid (TCA) cycle, nucleotide and hexosamine biosynthesis, redox homeostasis, and protein synthesis^3,4^. These pathways draw on a shared intracellular glutamine pool, yet the mechanisms that favor one use over another under nutrient limitation remain incompletely defined. Previous studies have largely focused on glutamine uptake or individual metabolic reactions, whether the machinery consuming glutamine for translation can itself alter the distribution of glutamine-derived resources is less clear.

Among the metabolic branches downstream of glutamine, polyamine biosynthesis represents an intriguing yet poorly understood route. Although spermine is conventionally synthesized from ornithine derived from arginine metabolism and the urea cycle^5,6^. Isotope tracing studies in PDAC indicate that glutamine can serve as a major carbon source for ornithine production^7^. The use of glutamine-derived precursors to produce spermine despite limited nutrient availability raises the possibility that this polyamine has regulatory roles beyond supporting cell growth. Indeed, spermine can be covalently attached to proteins through spermylation, a modification implicated in changes in protein function, localization, and interactions^8,9^. Whether spermylation allows spermine to influence other metabolic processes remains unclear.

Aminoacyl-tRNA synthetases (aaRSs) are positioned at the interface of nutrient metabolism and gene expression^10,11^. By coupling amino acids to cognate tRNAs, they determine the immediate supply of substrates for ribosomal elongation^12^. Recent studies further suggest that aaRS activity can influence translation of proteins enriched in particular amino acids and thereby contribute to metabolic adaptation in cancer^13,14^. Glutaminyl-tRNA charging becomes limiting during amino acid deprivation in some cellular settings^15^, but how metabolic signals regulate QARS aminoacylation when glutamine is needed for multiple competing processes remains poorly understood.

Here, we identify spermylation of QARS as a checkpoint in glutamine allocation. transglutaminase 2 (TGM2)-dependent spermylation enhances QARS aminoacylation and glutaminyl-tRNA charging, favoring glutamine incorporation into protein under nutrient stress. The resulting increase in charged tRNA also preferentially supports translation of glutamine codon rich transcripts, linking coding-sequence composition to metabolic reprogramming. These findings therefore connect a metabolite-derived covalent modification of the translation machinery to nutrient allocation and sequence-dependent translational selectivity.

## Results

### Spermine directs glutamine allocation to translation under nutrient stress

To investigate how glutamine use changes during pancreatic tumorigenesis, we first profiled metabolites in human PDAC and matched adjacent non-tumor tissues. Polyamines were prominent among the tumor-enriched metabolites, with increases in spermine and putrescine (Figure 1A). Targeted profiling confirmed increased spermine abundance in PDAC (Figure 1B). In *Kras*^LSL-G12D/+^ *Trp53*^R172H/+^ *Pdx1*-Cre (KPC) mice, spermine increased progressively from pancreatic intraepithelial neoplasia (PanIN) to invasive carcinoma, accompanied by increases in ornithine and putrescine and a decrease in spermidine (Figure 1C). These changes were consistent with increased spermine production and coincided with higher expression of the biosynthetic enzymes ODC1, AMD1, and SMS and lower expression of the catabolic enzyme PAOX. Clinically, higher SMS expression was associated with poorer prognosis (Figure 1D, Supplementary Figure 1A, B).

**Figure 1.**
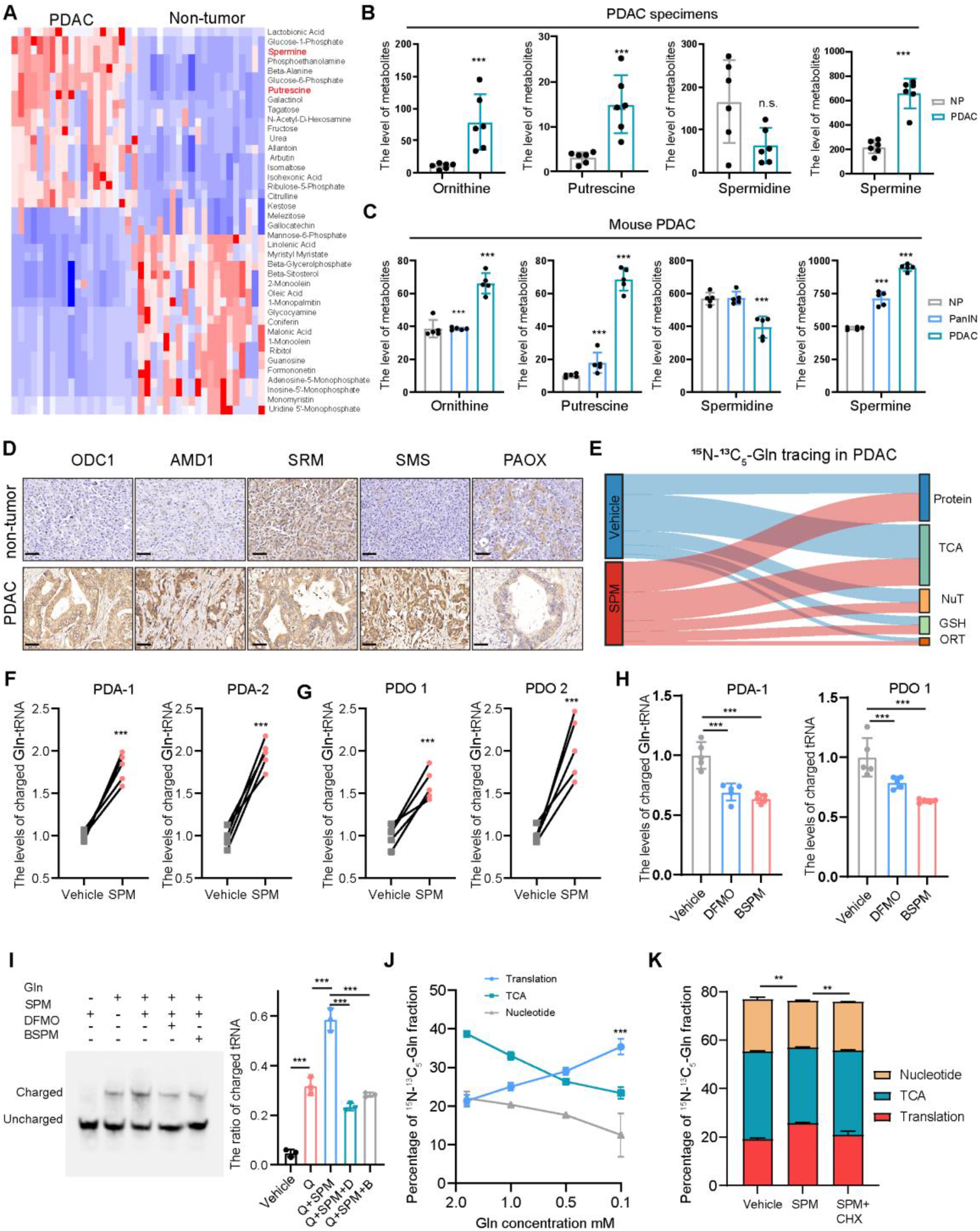
Spermine redistributes glutamine toward protein synthesis under nutrient stress. A. Heatmap of differential metabolites detected by untargeted liquid chromatography–mass spectrometry (LC–MS) in paired human pancreatic ductal adenocarcinoma (PDAC) and adjacent non-tumor tissues. Metabolite intensities were normalized to tissue mass. n = 20. **B**. Targeted isotope-dilution LC–MS/MS quantification of ornithine, putrescine, spermidine and spermine in paired human PDAC and adjacent non-tumor tissues. Metabolite abundances were normalized to tissue wet weight. n = 6. **C**. Targeted polyamine profiling across normal pancreas, PanIN lesions and PDAC in KPC mice. n = 5. **D**. Representative immunohistochemical staining for enzymes involved in spermine metabolism in human PDAC and adjacent non-tumor tissues. **E**. Pool-size-weighted relative allocation of label from ^13^C_5_,^15^N_2_-glutamine among protein-bound amino acids, tricarboxylic acid (TCA)-cycle intermediates and nucleotide pools in primary PDAC cells treated with vehicle or spermine (10 μM). Cells were incubated for 1 h in medium containing 1% dialyzed fetal bovine serum (FBS), 1 mM glucose and 0.5 mM labeled glutamine. **F-G**. Relative abundance of charged Gln-tRNA in primary PDAC cells (F) and patient-derived organoids (G) following spermine treatment. **H**. Charged Gln-tRNA abundance following inhibition of spermine synthesis with DFMO or BSPM in primary PDAC cells and PDOs. n = 3. **I**. Representative acid-urea northern blots resolving charged and uncharged tRNA^Gln^ after the indicated treatments; charging fractions were calculated as charged signal divided by total charged plus uncharged signal. **J**. Allocation of U-¹³C,¹ N -glutamine-derived label among protein, TCA-cycle and nucleotide pools across graded extracellular glutamine concentrations in the presence or absence of spermine. n = 5. **K**. Glutamine-derived label allocation in primary PDAC cells treated with vehicle, spermine or spermine plus cycloheximide (CHX), testing the dependence of spermine-induced redistribution on ongoing translation.

We next asked whether spermine changes the fate of intracellular glutamine rather than simply increasing its acquisition. Under nutrient limitation, spermine did not significantly alter total glutamine uptake (Supplementary Figure 1C). Nevertheless, stable-isotope tracing with ^13^C,^15^N-glutamine showed greater incorporation of glutamine-derived label into protein and lower labeling of TCA-cycle metabolites and nucleotide pools, labeling of glutathione and polyamines was largely unchanged in cells and tumors (Figure 1E, Supplementary Figure 1D, E). We refer to this redistribution of glutamine-derived labeling as glutamine allocation. To understand how spermine promotes the incorporation of glutamine into protein, we examined whether it regulates glutaminyl-tRNA charging, which directly links glutamine availability to translation. Spermine increased the abundance of charged tRNA^Gln^ in primary PDAC cells and patient-derived organoids (PDO), whereas inhibition of spermine biosynthesis reduced tRNA^Gln^ charging (Figure 1F–H). Northern blot analysis independently confirmed these changes in charged Gln-tRNA abundance (Figure 1I).

To test whether this redistribution becomes more pronounced during nutrient stress, we varied extracellular glutamine concentrations. As glutamine availability decreased, spermine preferentially maintained protein synthesis while reducing glutamine-derived labeling of competing pathways (Figure 1J). Moreover, low-dose cycloheximide largely prevented the spermine-associated redistribution of glutamine-derived label (Figure 1K), consistent with translation acting as a major competing sink for glutamine. Together, these results indicate that spermine favors glutamine allocation to protein synthesis when nutrients are scarce.

### QARS spermylation enhances protein translation

We next investigated how spermine alters glutamine allocation. Because spermine has been implicated in reactive oxygen species signaling, autophagy, and JAK signaling^16–18^, we first tested whether these pathways contributed to the observed metabolic response. Their pharmacologic inhibition did not prevent spermine-associated glutamine redistribution (Figure 2A). By contrast, GK921, an inhibitor of TGM2, reversed this response (Figure 2A), suggesting a role for covalent spermine attachment to proteins. Using the prime adenosine-retaining aminoacyl-tRNA isolation (PARTI)^19^ assay, we found that spermine increased Gln-tRNA charging in primary PDAC cells, whereas GK921 abolished this increase (Figure 2B). To identify candidate covalently modified proteins, we applied an alkyne-tagged spermine probe and enriched labeled proteins for mass spectrometry. QARS emerged as a prominent candidate alongside other recovered proteins (Supplementary Figure 2A; Supplementary Table 1). QARS enrichment persisted after high-salt or 8 M urea washing, consistent with a stable, potentially covalent association (Figure 2C). Immunofluorescence showed co-localization of QARS with alkyne-spermine in primary PDAC cells and with endogenous spermine in clinical PDAC specimens (Figure 2D). In a complementary *in vitro* transamidation assay, active TGM2 increased incorporation of the fluorescent substrate monodansylcadaverine (MDC) into QARS, whereas TGM2 inhibition reduced incorporation (Figure 2E).

**Figure 2.**
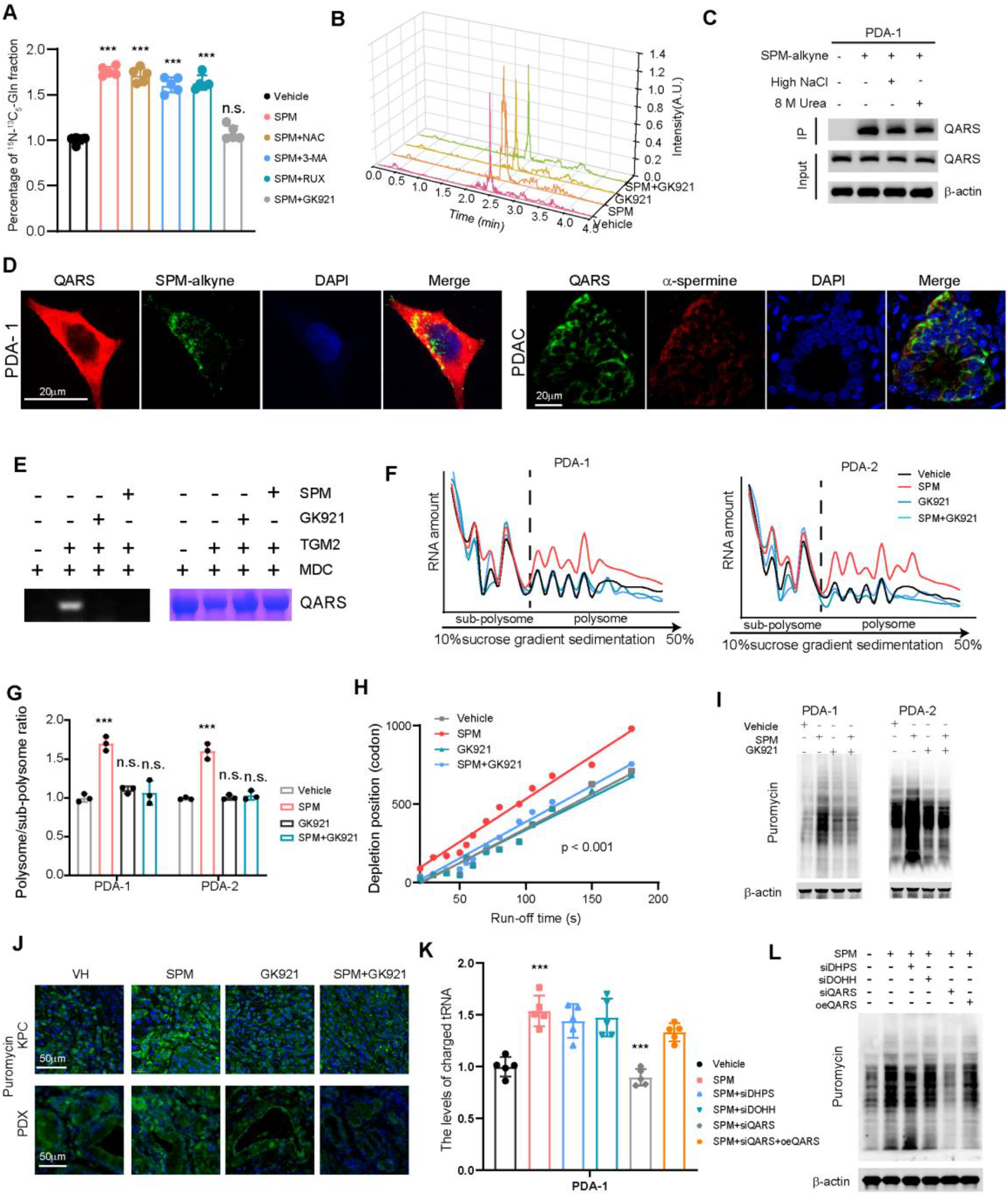
Spermylation of QARS enhances protein translational efficiency. **A**. Protein-bound ^15^N amino acids in primary cells were treated with spermine in combination with the ROS inhibitor N-acetylcysteine (NAC), the autophagy inhibitor 3-methyladenine (3-MA), the JAK pathway inhibitor ruxolitinib (RUX) or the transglutaminase 2 (TGM2) inhibitor GK921 (TGMi), as specified. n = 5.**B**. Relative abundance of charged Gln-tRNA in primary PDAC cells treated with vehicle, 10 μM spermine, GK921 or spermine plus GK921. n = 5. **C**. Anti-QARS immunoblot of proteins enriched by alkyne-spermine labeling, click chemistry and streptavidin pull-down, including stringent high-salt and 8 M urea washing conditions. **D**. Immunofluorescence staining of QARS together with alkyne spermine in PDAC cells or spermine in pancreatic cancer specimens. **E**. *In vitro* TGM2-mediated QARS transamidation assays performed in the presence or absence of GK921 or excess spermine. **F-G**. Representative polysome profiles (F) and quantification of the polysome-to-subpolysome ratio (G); n = 3. **H**. Ribosome run-off following harringtonine addition (2 μg /ml; 0, 1, 2, 3, 4 and 5 min), measured as the remaining polysome fraction normalized to the 0-min sample in cells treated with vehicle, spermine, GK921 or the combination. Sample size and replication: n = 3 independent biological experiments **I**. Representative puromycin incorporation assays and corresponding quantification of nascent protein synthesis in PDA-1 and PDA-2 cells following the indicated treatments. Puromycin signal was integrated across the blot lane and normalized to total-protein staining **J**. In vivo puromycin incorporation (20 mg/kg, intraperitoneal injection; tumors collected after 30 min) in KPC and patient-derived xenograft (PDX) tumors treated with spermine, with or without GK921. **K**. Charged Gln-tRNA abundance in PDAC cells after siRNA-mediated depletion of deoxyhypusine synthase (DHPS), deoxyhypusine hydroxylase (DOHH) or QARS, and after QARS re-expression in the indicated rescue groups. n = 5. **L**. Puromycin incorporation in PDA-1 cells after depletion of DHPS, DOHH or QARS, or after QARS re-expression, as specified.

We then tested whether spermine-dependent spermylation increases protein synthesis. Spermine increased the polysome-to-subpolysome ratio and accelerated apparent ribosome run-off, whereas GK921 counteracted the run-off response (Figure 2F–H; Supplementary Figure 2B). Puromycin incorporation likewise supported increased nascent protein production following spermine treatment and reduced translation after interference with spermine synthesis or spermylation (Figure 2I; Supplementary Figure 2C). Similar changes in puromycin incorporation were observed in KPC and patient-derived xenograft tumors (Figure 2J).

Because polyamines also influence translation through eIF5A hypusination^20^, we examined the contribution of this established pathway. Depletion of deoxyhypusine synthase (DHPS) or deoxyhypusine hydroxylase (DOHH) had little effect on spermine-induced Gln-tRNA charging or polysome accumulation (Figure 2K, Supplementary Figure 2D-E). In contrast, QARS knockdown markedly reduced charged tRNA^Gln^ abundance and the P/S ratio despite spermine supplementation. Ectopic QARS expression restored charging and polysome formation in spermine-treated cells (Figure 2K, Supplementary Figure 2E). Puromycin incorporation provided parallel evidence: inhibition of DHPS or DOHH had little effect on the spermine response, whereas QARS depletion largely eliminated the increase in nascent protein synthesis (Figure 2L). QARS is a major mediator of the translational response to spermine in PDAC models.

### Spermylation at QARS Q216 promotes aminoacylation and glutamine allocation to translation

We next investigated how spermylation regulates QARS itself. Mass spectrometry of in vitro spermylated recombinant QARS identified modified glutamine residues at Q41, Q86 and Q216 (Figure 3A; Supplementary Figure 3A). Q216A retained basal aminoacylation activity similar to that of wild-type QARS but failed to show the spermine-dependent increase in activity (Figure 3B; Supplementary Figure 3B-C). Molecular dynamics simulations suggested that Q216 spermylation reduced fluctuations in residues 50-90 and 90-110 upon tRNA binding (Supplementary Figure 3D). The modified QARS-tRNA complex also reached a stable conformation earlier and exhibited a lower maximal root-mean-square deviation than the unmodified complex (Figure 3C). These calculations do not establish the physical binding mechanism, but are consistent with altered substrate-complex dynamics. Independent *in vitro* aminoacylation assays confirmed that spermylation enhanced QARS catalytic activity and that this enhancement was prevented by TGM2 inhibition or Q216 mutation (Figure 3D-E).

**Figure 3.**
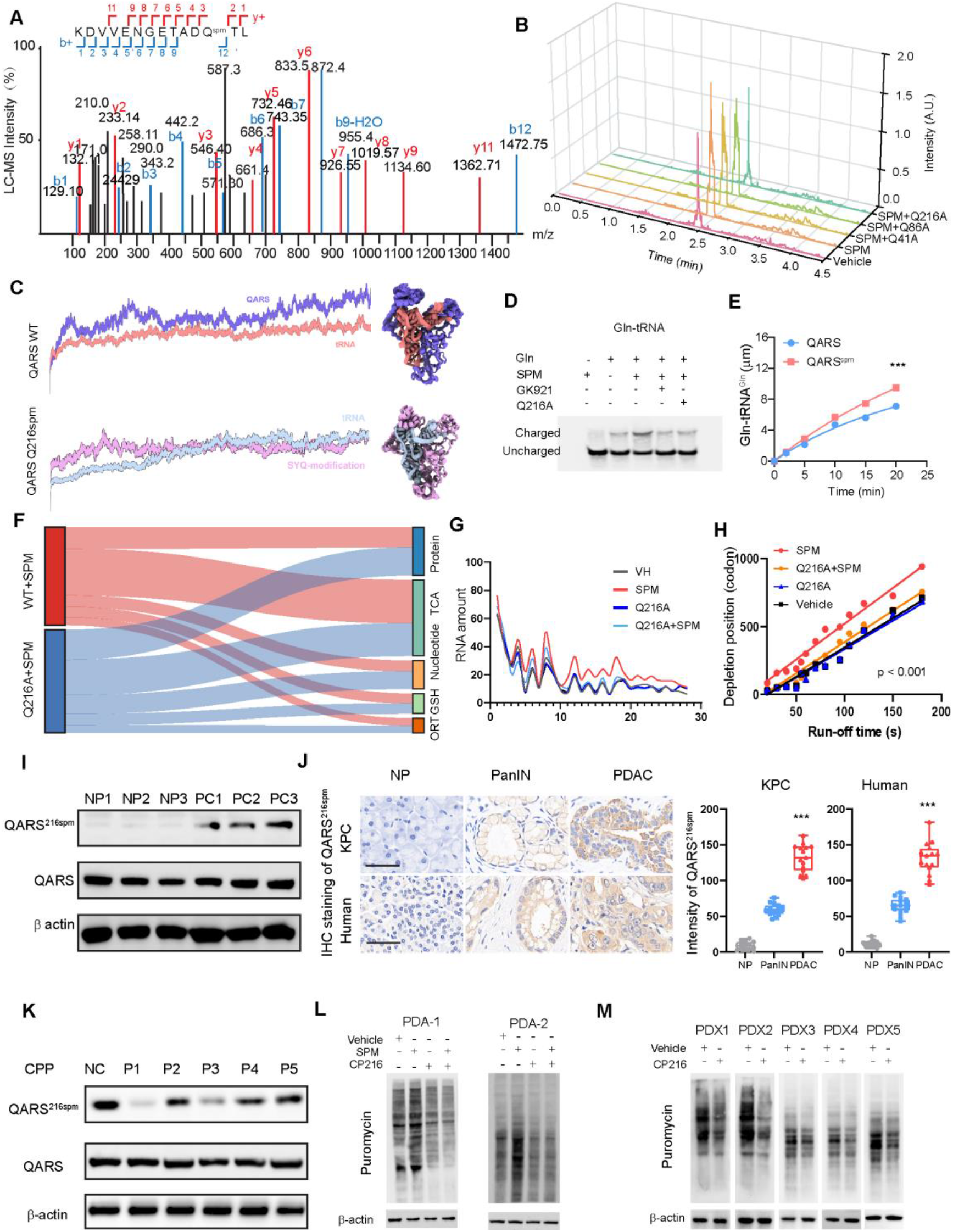
QARS Q216 spermylation enhances aminoacylation and channels glutamine toward translation. **A.** LC-MS/MS identification of spermylated glutamine residues in QARS residues Q216. **B**. Relative levels of charged Gln-tRNA in QARS-WT, Q41A, Q86A and Q216A PDAC cells following spermine treatment. n = 3 independent biological experiments. **C**. Molecular dynamics analysis of Gln-tRNA bound to unmodified or Q216-spermylated QARS. **D**. Northern blot analysis of charged and uncharged Gln-tRNA after pharmacologic inhibition of spermylation or Q216A mutation. **E**. Time course of Gln-tRNA^Gln^ formation catalyzed by recombinant unmodified or spermylated QARS in vitro (0, 1, 2, 4 and 6 min), with initial rates calculated from the linear phase. **F**. ^13^C_5_,^15^N_2_-glutamine-derived label distribution among protein-bound amino acids, TCA-cycle metabolites and nucleotides in in QARS-WT and Q216A knock-in primary PDAC cells under nutrient-limited conditions (1% dialyzed FBS, 1 mM glucose and 0.5 mM glutamine; 1 h). **G-H**. Polysome-to-subpolysome ratio (G) and ribosome run-off kinetics (H) in QARS-WT and Q216A primary PDAC cells following spermine treatment. **I**. Immunoblot detection of QARS Q216 spermylation and total QARS in patient-matched PDAC and adjacent non-tumor tissues. **J**. Immunohistochemical staining for QARS Q216 spermylation in non-tumor pancreas, PanIN lesions and invasive PDAC. Quantification is based on staining intensity. n=10. **K**. Immunoblot validation of CP216-mediated blockade of QARS Q216 spermylation in PDA-1 cells. **L**. Puromycin incorporation in primary PDAC cells treated with vehicle, spermine, CP216 or spermine plus CP216. **M**. Puromycin incorporation in patient-derived xenograft tumors treated with control peptide or CP216.

To determine whether this molecular change affects nutrient utilization, we compared QARS-WT and Q216A PDAC cells. Q216A decreased glutamine-derived labeling of protein while increasing labeling of TCA-cycle and nucleotide pools (Figure 3F; Supplementary Figure 3E). Q216A also prevented spermine-induced increases in the polysome-to-subpolysome ratio and apparent ribosome run-off rate (Figure 3G-H). To examine Q216 spermylation in tissues, we developed a modification-specific antibody against QARS Q216 spermylation (anti-QARS^Q216spm^) and validated its specificity and sensitivity (Supplementary Figure 3F). Immunoblotting detected QARS^Q216spm^ in PDAC samples but little signal in matched adjacent non-tumor tissues (Figure 3I). Immunohistochemistry further showed progressively stronger staining from adjacent non-tumor tissue to PanIN lesions and invasive PDAC (Figure 3J), linking the modification to disease progression.

To block QARS spermylation acutely, we designed cell-penetrating peptides (CPPs) containing sequences surrounding Q216 to compete with modification of the endogenous site. Screening in primary PDAC cells identified P1, hereafter CP216, as the most effective peptide. CP216 reduced Q216 spermylation with little change in total QARS abundance (Figure 3K). CP216 also prevented spermine-induced Gln-tRNA charging and decreased nascent protein synthesis in primary PDAC cells (Figure 3L; Supplementary Figure 3G), with reduced translation observed in PDX tumors (Figure 3M). These data provide an acute perturbation that broadly phenocopies the Q216A genetic intervention.

### QARS spermylation preferentially sustains translation of glutamine codon rich transcripts

To determine whether QARS spermylation affects translation broadly or favors particular transcripts, we performed ribosome profiling (Ribo-seq) in PDAC cells treated with scrambled CPP or CP216 (Supplementary Figure 4A; Supplementary Table 2). Transcripts with higher TE when QARS spermylation was intact were enriched for functions in nucleotide and carbohydrate metabolism, energy production, and proliferation (Figure 4A-B). Among the candidates were UPP1, UCK2, and CAD, along with the growth regulator MYC (Figure 4C). Ribosome occupancy on these transcripts increased after spermine treatment, as confirmed in primary PDAC cells (Figure 4D), whereas mRNA levels were largely unchanged (Supplementary Figure 4B), indicating regulation primarily at the level of translation. Immunoblotting corroborated increased UPP1, UCK2, and MYC protein abundance after spermine treatment. Inhibition of spermine biosynthesis or TGM2 reduced their expression (Supplementary Figure 4C-D), and Q216A prevented their induction by spermine (Figure 4E). In KPC tumors, CP216 likewise reduced QARS Q216 spermylation and the abundance of UPP1, UCK2, and MYC (Figure 4F).

**Figure 4.**
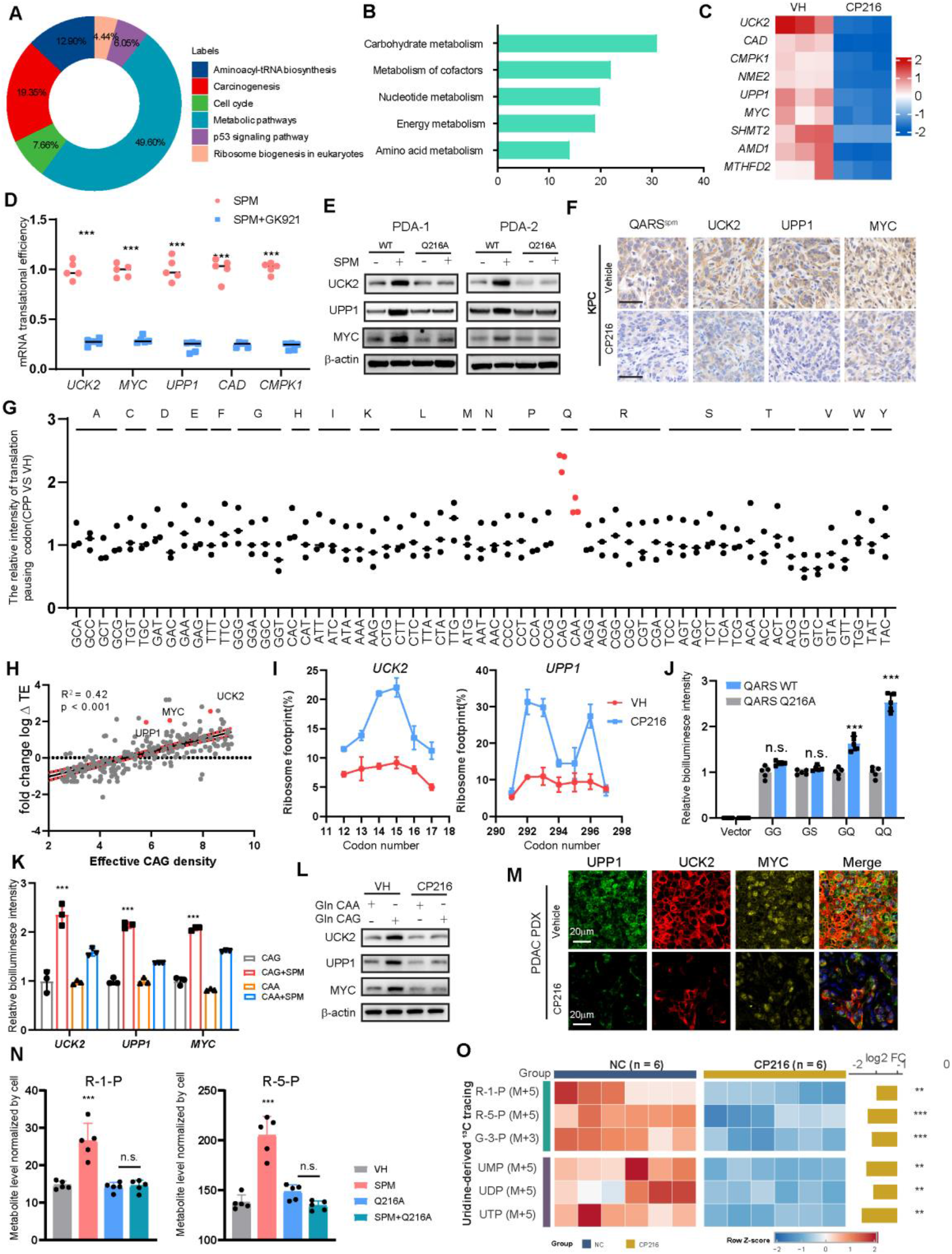
QARS spermylation establishes glutamine-codon-dependent translational prioritization. **A**.Circular visualization of Kyoto Encyclopedia of Genes and Genomes (KEGG) pathway enrichment for transcripts with altered translational efficiency following treatment with CP216 versus scrambled control peptide, determined from ribosome profiling with matched total RNA-seq. **B**. Enrichment of metabolic KEGG pathways among transcripts showing QARS-spermylation-dependent changes in translational efficiency. **C**.Heatmap of representative metabolism-related transcripts with differential translational efficiency following CP216 treatment. **D**. Validation of selected QARS-spermylation-dependent transcripts in primary PDAC cells following spermine treatment. n=5. **E**. Representative immunoblots and quantification of UPP1, UCK2 and MYC abundance in QARS-WT and Q216A PDAC cells after vehicle or spermine treatment. **F**. Immunohistochemistry for QARS Q216 spermylation, UPP1, UCK2 and MYC in KPC tumors treated with scrambled cell-penetrating peptide (CPP) or CP216. **G**. A-site codon occupancy analysis showing changes in ribosome pausing at individual codons following CP216-mediated blockade of QARS spermylation. n=3. **H**. Association between effective CAG density and QARS-spermylation-dependent translational efficiency. **I**. Codon-resolved ribosome footprint profiles across UCK2 and UPP1 following scrambled CPP or CP216 treatment. n=3. **J**. Firefly luciferase reporter activity from constructs containing graded numbers of glutamine codons in QARS-WT and Q216A cells after the indicated treatments, normalized to a cotransfected Renilla reporter. n=5. **K-L**. Functional and immunoblot analyses of synonymous CAG-versus CAA-recoded UCK2, UPP1 and MYC constructs. M. Multiplex immunofluorescence of UPP1, UCK2 and MYC in PDX tumors treated with scrambled CPP or CP216. **N**. Relative levels of ribose-1-phosphate (R-1-P) and ribose-5-phosphate (R-5-P) in PDAC cells following manipulation of spermine/QARS Q216 spermylation. n=5. **O**. U-13C5-uridine tracing in KPC tumors showing incorporation of uridine-derived carbon into downstream metabolic intermediates after scrambled CPP or CP216 treatment. n=6.

We next examined the coding-sequence features associated with sensitivity to QARS spermylation. Codon-resolved Ribo-seq analysis showed increased ribosome pausing at glutamine codons after CP216 treatment, with smaller changes at other codons (Figure 4G), consistent with reduced charged tRNA^Gln^ availability. Transcripts whose translation depended more strongly on QARS spermylation contained more glutamine codons (Figure 4H). CAG was the predominant glutamine codon in UCK2 and several other responsive transcripts (Supplementary Figure 4E-F). Correspondingly, ribosome footprints accumulated at glutamine codons within UPP1 and UCK2 after CP216 treatment (Figure 4I), indicating local constraints on elongation.

To test sequence dependence more directly, we constructed luciferase reporters containing increasing numbers of glutamine codons. Greater glutamine-codon content conferred stronger dependence on QARS spermylation, whereas Q216A attenuated this relationship (Figure 4J, Supplementary Figure 4G). We then generated synonymous CAA-enriched and CAG-enriched versions of UPP1, UCK2 and MYC, preserving their amino acid sequences. Spermine preferentially increased protein expression from the CAG-enriched constructs relative to matched CAA-enriched constructs, and CP216 largely eliminated this difference (Figure 4K-L). Consistent with the cell-based findings, multiplex immunofluorescence showed reduced UPP1, UCK2, and MYC abundance in CP216-treated PDX tumors (Figure 4M).

We then assessed the metabolic consequences of preferentially translating uridine-metabolism enzymes. In primary PDAC cells, spermine increased ribose-1-phosphate (R-1-P) and ribose-5-phosphate (R-5-P) abundance, an effect absent with Q216A (Figure 4N). We further performed U-^13^C_5_-uridine tracing in KPC tumors. CP216 reduced incorporation of uridine-derived carbon into R-1-P, R-5-P, and pyrimidine nucleotide pools (Figure 4O), supporting a role for QARS spermylation in promoting uridine utilization *in vivo*.

Together, these findings indicate that QARS spermylation preferentially sustains translation of glutamine-codon-rich transcripts by enhancing Gln-tRNA charging, an effect more pronounced in CAG-enriched than in matched CAA-recoded constructs. This translational response supports uridine-metabolic enzymes and may reduce reliance on glutamine-intensive de novo nucleotide synthesis.

### QARS Q216 spermylation sustains PDAC progression *in vivo*

We next assessed the biological importance of this mechanism in PDAC. In orthotopic tumors generated from KPC-derived cells, Q216A abolished the spermine-associated increase in tumor growth and prolonged survival (Figure 5A–C). Puromycin incorporation showed lower nascent protein synthesis in Q216A tumors and loss of the translational response to spermine (Figure 5D). To examine glutamine allocation directly, we performed U-^13^C_5_,^15^N_2_-glutamine tracing in orthotopic tumors. Spermine increased protein-bound ^15^N labeling in wild-type tumors, whereas Q216A prevented this increase (Figure 5E). In contrast, Q216A increased incorporation of glutamine-derived label into α-ketoglutarate and nucleotide pools, including UMP isotopologues (Figure 5F-G), supporting redistribution among competing metabolic pools *in vivo*.

**Figure 5.**
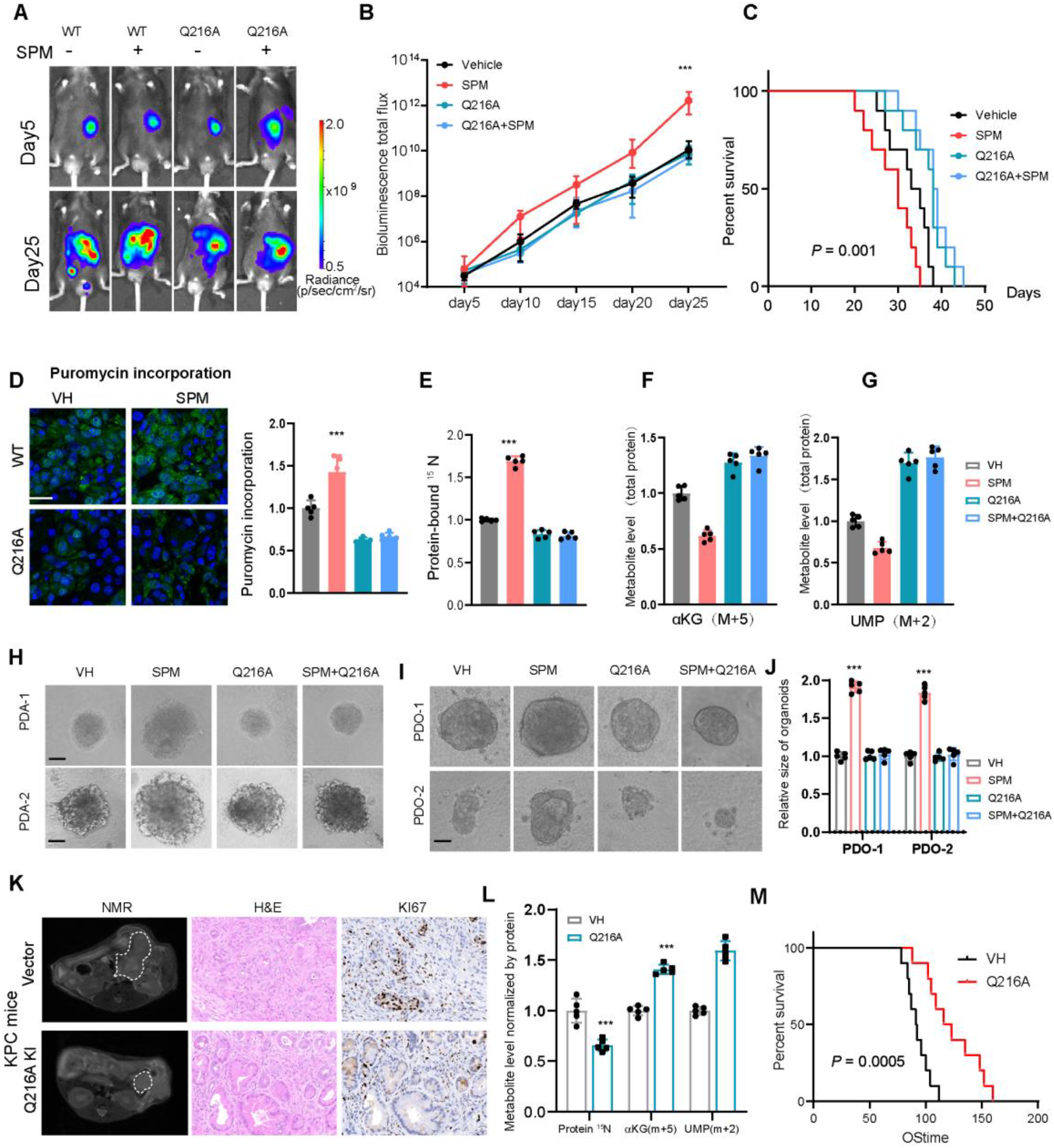
QARS Q216 spermylation promotes PDAC progression in vivo. **A**. Representative serial bioluminescence images of orthotopic tumors established from QARS-WT or Q216A KPC-derived PDAC cells in mice treated with vehicle or spermine. n = 5. **B**. Longitudinal quantification of tumor bioluminescence (total photon flux) for the groups shown in orthotopic tumors generated from QARS-WT or Q216A KPC-derived PDAC cells with or without spermine. n = 5. **C**. Kaplan–Meier survival curves of mice bearing orthotopic QARS-WT or Q216A tumors treated with vehicle or spermine. n=10. D. *In vivo* puromycin incorporation in orthotopic QARS-WT or Q216A tumors following vehicle or spermine treatment. n=3. E-G. ^13^C_5_,^15^N_2_-glutamine tracing in orthotopic tumors showing protein-bound ^15^N (E), α-ketoglutarate M+5 (F), and glutamine-derived labeling of UMP isotopologues (G). **H**. Three-dimensional growth of primary PDAC cells expressing QARS-WT or Q216A following vehicle or spermine treatment. n = 3. **I-J**. Representative patient-derived organoids (I) and quantification of organoid area (J) under the indicated conditions. **K**. Representative magnetic resonance imaging (MRI), hematoxylin-and-eosin (H&E) and Ki67 staining of tumors in autochthonous KPC mice following adeno-associated virus (AAV)-mediated QARS-WT or Q216A editing. n = 5. **L**. Protein-bound ^15^N, α-ketoglutarate M+5 and glutamine-derived labeling of UMP isotopologues labeling in autochthonous KPC tumors carrying AAV-mediated QARS-WT or Q216A edits following^13^C_5_,^15^N_2_-glutamine tracing. n = 5. **M**. Kaplan–Meier overall survival of KPC mice following AAV-mediated QARS-WT or Q216A editing. n = 10.

We next extended these observations to additional disease-relevant models. Q216A impaired the spermine-associated increase in spheroid growth in primary PDAC cultures (Figure 5H); patient-derived organoids similarly lost their growth response when Q216 spermylation was blocked (Figure 5I-J). For genetic assessment in an autochthonous setting, we used AAV-mediated CRISPR delivery to generate QARS-WT and Q216A KPC cohorts (Supplementary Figure 5A). MRI and histopathology showed lower tumor burden and less advanced lesions in Q216A KPC mice, accompanied by reduced Ki67 staining (Figure 5K, Supplementary Figure 5B-D). Tumors from Q216A mice also exhibited lower protein-bound ^15^N labeling, reciprocal increases in glutamine-derived α-ketoglutarate and UMP labeling (Figure 5L), and prolonged survival (Figure 5M). These findings support a role for Q216 spermylation in PDAC progression and glutamine allocation *in vivo*.

### Peptide-mediated inhibition of QARS spermylation suppresses PDAC progression

We next examined the therapeutic potential of inhibiting QARS Q216 spermylation. After a single intraperitoneal injection of CP216 (10 mg/kg), plasma concentrations showed an apparent terminal half-life of 3.75 h and remained measurable for 8 h (Supplementary Figure 6A). Repeated administration was well tolerated under the tested regimen, with no significant changes in body weight, serum markers of liver or kidney function, or major-organ histology (Supplementary Figure 6B-D). In established PDA-1 subcutaneous xenografts, four weeks of CP216 treatment reduced tumor growth and endpoint weight compared with scrambled CPP controls (Figure 6A-B, Supplementary Figure 6E). Immunohistochemistry confirmed lower QARS Q216 spermylation and proliferation in treated tumors (Figure 6C). CP216 similarly suppressed growth across multiple PDX models, with reduced QARS Q216 spermylation confirming target engagement (Figure 6D, Supplementary Figure 6F). To test whether competing metabolic demands influence the response to QARS blockade, we treated mice bearing orthotopic PDAC tumors with CP216 plus dimethyl-α-ketoglutarate and ribonucleosides (KG&R), which supply metabolites downstream of the TCA cycle and nucleotide biosynthesis.

**Figure 6.**
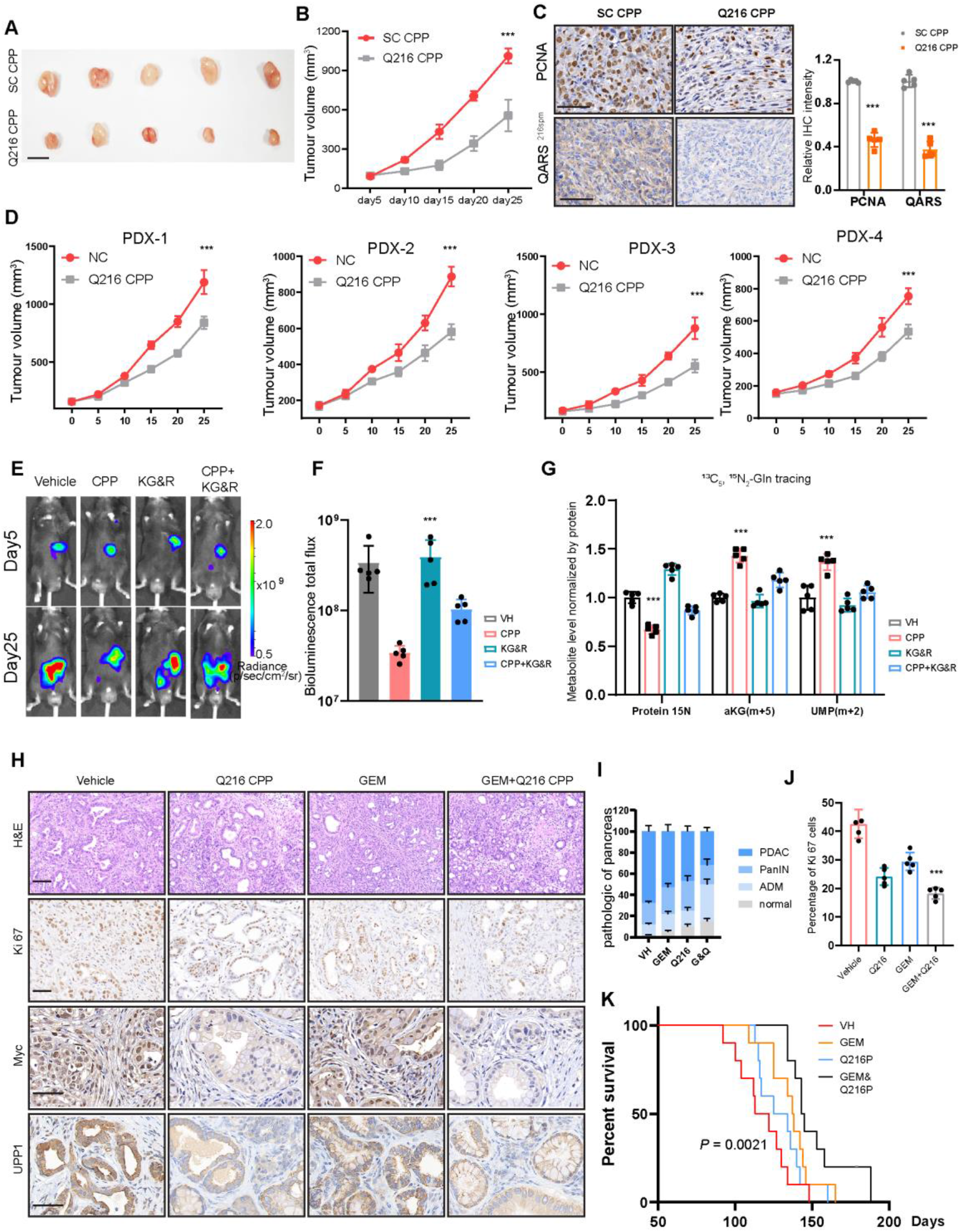
Targeting QARS spermylation suppresses PDAC progression in vivo. **A**. Representative excised tumors from PDA-1 subcutaneous xenografts treated with scrambled control CPP (SC-CPP) or CP216 (10 mg/kg, intraperitoneally, every 3 days for 4 weeks). n = 5. **B**. Longitudinal tumor volumes for the PDA-1 subcutaneous xenografts, measured by caliper and calculated as length × width²/2, with mouse as the repeated-measures unit. n = 5. **C**. Representative IHC staining and quantification of PCNA and QARS Q216 spermylation in SC-CPP- and CP216-treated tumors. n = 5.**D**. Tumor growth in four independent PDAC patient-derived xenograft models treated with control peptide or CP216. n = 5. **E**. Representative longitudinal bioluminescence images of orthotopic PDAC-bearing mice treated with vehicle, CP216, KG&R, or CP216 plus KG&R. n = 5. **F**. Quantification of endpoint bioluminescence in the groups shown in E. **G**. ^13^C_5_,^15^N_2_-glutamine-derived labeling of protein-bound amino acids, TCA-cycle metabolites and nucleotide pools. n = 5. **H**. Representative H&E and immunohistochemical staining for Ki67, MYC and UPP1 in KPC tumors treated with vehicle, CP216, gemcitabine (GEM; 50 mg/kg, intraperitoneal) or CP216 plus GEM **I**. Histological quantification of normal pancreas, acinar-to-ductal metaplasia (ADM), PanIN and invasive PDAC regions in the four KPC treatment groups. n = 5. **J**. Quantification of Ki67-positive cells in KPC tumors. n = 5.**K**. Kaplan-Meier survival analysis of KPC mice treated with vehicle, GEM, CP216, or GEM plus CP216. n = 10.

KG&R partially rescued growth of CP216-treated tumors, as measured by serial bioluminescence imaging and endpoint tumor burden (Figure 6E-F). Isotope tracing showed lower glutamine-derived labeling of TCA-cycle and nucleotide pools alongside partial restoration of labeling in protein (Figure 6G). These results suggest that competition among glutamine-consuming pathways, in addition to translational capacity, influences the response to QARS spermylation blockade.

Given the connection between gemcitabine action and nucleotide metabolism, we also tested CP216 together with gemcitabine in KPC mice. Combined treatment reduced histologic PDAC burden more than either agent alone (Figure 6H-I), accompanied by fewer Ki67-positive cells (Figure 6J) and lower MYC and UPP1 staining. In autochthonous KPC mice, CP216 prolonged median survival relative to vehicle (Figure 6K). These results provide proof of principle for therapeutic inhibition of QARS spermylation and suggest that CP216 can enhance gemcitabine activity in the models examined.

### QARS Q216 spermylation correlates with aggressive disease and poor outcomes

Finally, we examined the clinical relevance of QARS Q216 spermylation. Immunohistochemistry with the QARS Q216spm-specific antibody showed stronger staining in tumors than in matched nonmalignant tissues across PDAC, colorectal cancer (CRC), stomach adenocarcinoma (STAD), hepatocellular carcinoma (HCC), and lung adenocarcinoma (LUAD) cohorts (Figure 7A-B; Supplementary Figure 7A-B). In a tissue microarray of 150 clinically annotated PDAC specimens, we classified cases as QARS Q216spm-high or -low using the cohort median (Figure 7C-D). Higher QARS Q216spm staining was associated with larger tumors, more advanced TNM stage, and more frequent chemotherapy resistance (Figure 7E-G). Patients with higher staining also had shorter overall survival (Figure 7H), and associations with adverse outcomes were observed in the additional tumor cohorts examined (Supplementary Figure 7C). Together, the clinical and mechanistic results support a model in which spermine-dependent QARS spermylation sustains translation of glutamine-codon-rich metabolic programs and facilitates tumor growth under nutrient stress (Figure 7I).

**Figure 7.**
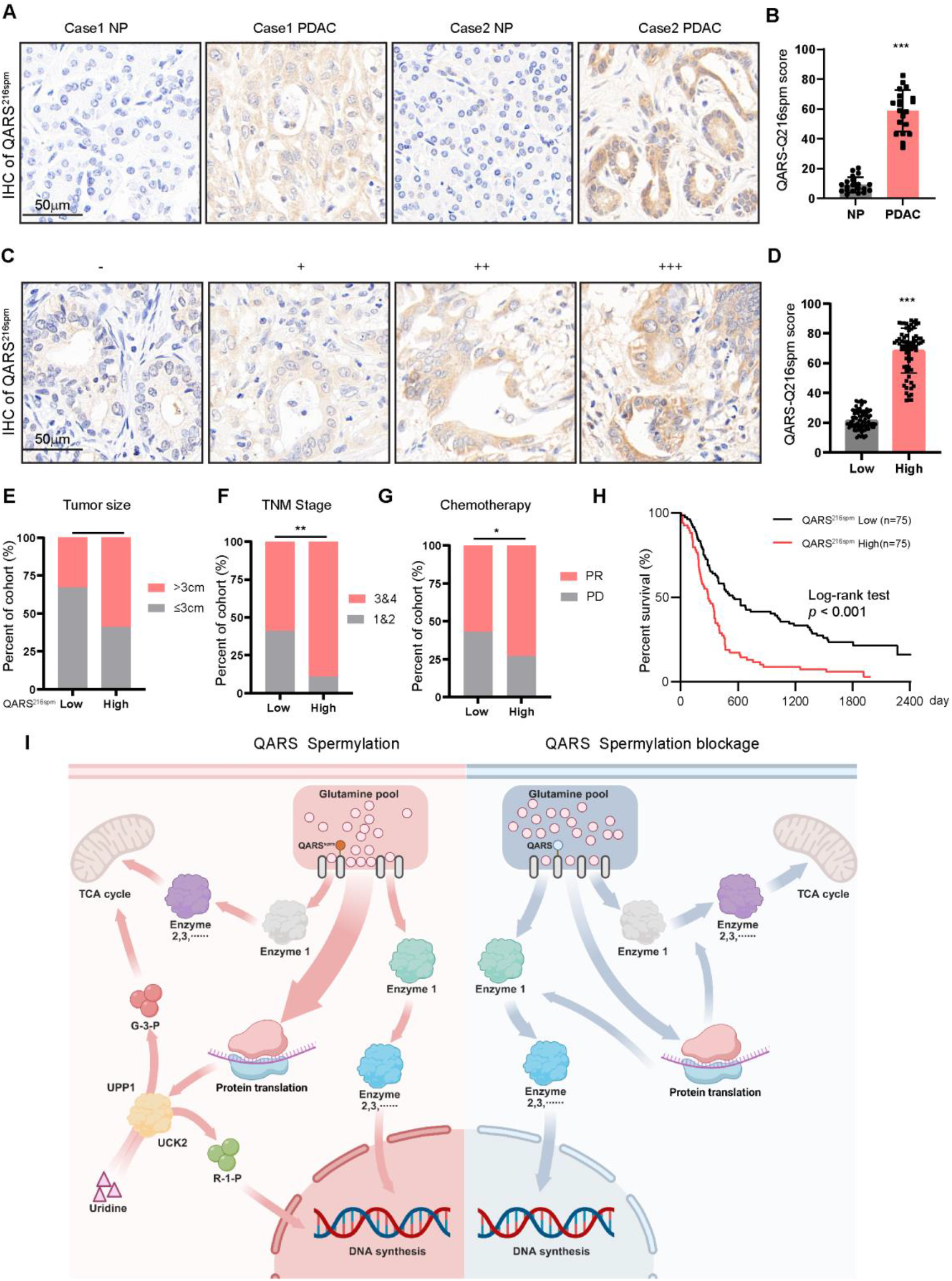
QARS Q216 spermylation is associated with aggressive PDAC and poor outcome. **A**. Representative immunohistochemical staining of QARS Q216 spermylation in PDAC and paired adjacent non-tumor tissues. Scale bar, 50 μm. **B**. Quantification of QARS Q216 spermylation scores in paired PDAC and adjacent non-tumor tissues (n = 20 pairs). **C**. Representative immunohistochemical staining illustrating negative, weak, moderate, and strong QARS Q216 spermylation in the PDAC tissue microarray. Scale bar, 50 μm. **D**. Distribution of QARS Q216 spermylation scores in the PDAC cohort (n = 150), dichotomized at the cohort median into low and high groups. **E-G**. Associations of QARS Q216 spermylation status with tumor size (E), TNM stage (F) and chemotherapy response (G). **H**. Kaplan–Meier overall-survival analysis of patients with QARS Q216spm-low (n = 75) or QARS Q216spm-high (n = 75) tumors. **I**. Working model in which QARS Q216 spermylation enhances Gln-tRNA charging and preferentially sustains translation of glutamine-codon-rich metabolic transcripts, promoting uridine utilization, metabolic adaptation and tumor growth.

## Discussion

Nutrient availability has traditionally been viewed as an upstream determinant of biosynthesis^21^, with translation regarded primarily as a downstream consumer of metabolic substrates^22,23^. Our findings reveal a reciprocal relationship in which regulation of the translation machinery can influence how a limiting nutrient is utilized. In PDAC, spermine-dependent QARS spermylation enhances glutamine incorporation into protein without increasing glutamine uptake, accompanied by reduced relative labeling of TCA-cycle metabolites and nucleotides. These observations identify translation as an active participant in metabolic resource allocation rather than simply an endpoint of nutrient supply.

The molecular basis of this regulation involves covalent modification of an enzyme directly responsible for incorporating glutamine into protein. Transglutaminase-dependent polyamination has previously been described for several protein substrates^24,25^. Here, we identify QARS Q216 as a functionally important spermylation site. Q216 spermylation enhances QARS aminoacylation and Gln-tRNA charging, whereas the Q216A mutation preserves basal catalytic activity while preventing spermine-dependent enhancement. Thus, spermylation modulates the canonical function of QARS rather than conferring an unrelated activity^26^, providing a mechanism through which polyamine abundance influences translational demand for glutamine. Importantly, our isotope-tracing measurements describe the relative distribution of glutamine-derived label rather than absolute, mass-balanced metabolic flux, and should be interpreted within this framework.

The consequences of QARS spermylation extend beyond an overall increase in protein synthesis. Ribosome profiling revealed glutamine-codon-associated pausing upon spermylation blockade, whereas transcripts enriched in glutamine codons showed greater dependence on the modification. Enhanced Gln-tRNA charging may therefore preferentially relieve elongation constraints on transcripts with high glutamine-codon demand^12^. Notably, synonymous recoding revealed stronger translational responses in CAG-enriched constructs than in their CAA-recoded counterparts, despite identical encoded proteins. Although the underlying mechanism remains unresolved, these findings suggest that synonymous codon usage can influence translational sensitivity to tRNA availability, potentially through differences in the abundance, charging efficiency, or decoding properties of the two mammalian tRNA^Gln^ isoacceptors that recognize CAG and CAA codons ^27^. QARS spermylation increases the availability of charged tRNA^Gln^, which may differentially affect transcripts depending on their glutamine codon composition and the properties of corresponding tRNA^Gln^ isoacceptors.

An important consequence of this translational selectivity is the preferential expression of enzymes that facilitate alternative nutrient utilization. Among these, UPP1 and UCK2 provide distinct routes for uridine metabolism^28,29^. UPP1 catalyzes uridine phosphorolysis to generate ribose-1-phosphate, allowing ribose-derived carbon to contribute to central carbon metabolism, including pathways connected to the TCA cycle. UCK2, in contrast, supports pyrimidine salvage by directly phosphorylating uridine to UMP. Our isotope-tracing experiments demonstrate that QARS spermylation increases the incorporation of uridine-derived carbon into ribose phosphates and nucleotide pools. Although increased uridine contribution to TCA-cycle intermediates remains to be established in our models, these observations suggest that QARS-dependent translation promotes the utilization of metabolic substrates beyond glutamine.

This metabolic adaptation may provide an explanation for the preferential translation of uridine-metabolic enzymes under nutrient stress. By increasing nucleotide salvage and potentially providing alternative carbon inputs for central metabolism, enhanced uridine utilization could alleviate some of the competing demands placed on glutamine, thereby favoring its incorporation into protein. Such a mechanism would allow tumor cells to sustain protein synthesis while making more effective use of available metabolic resources. Consistent with this interpretation, supplementation with α-ketoglutarate and ribonucleosides partially restored protein labeling and tumor growth following CP216-mediated spermylation blockade. These results support a model in which translational prioritization and alternative nutrient utilization reinforce one another to facilitate adaptation to nutrient scarcity. Nevertheless, the extent to which UPP1- and UCK2-dependent uridine metabolism directly spares glutamine remains to be determined through genetic perturbation and quantitative metabolic flux analysis.

The enrichment of QARS Q216 spermylation during PDAC progression also suggests a potential therapeutic opportunity. Direct inhibition of glutamine metabolism can affect normal tissues because of their dependence on this nutrient^30,31^. Targeting a tumor-enriched regulatory modification may therefore offer an alternative to broadly suppressing glutamine utilization. Genetic disruption of Q216 spermylation reduced tumor progression, while CP216 suppressed tumor growth across xenograft, PDX, and autochthonous models and enhanced the antitumor activity of gemcitabine. These findings provide proof of principle for peptide-mediated inhibition QARS spermylation, although its therapeutic selectivity and safety require further investigation.

Several questions remain. The structural basis by which Q216 spermylation enhances QARS aminoacylation and the mechanisms underlying the distinct translational responses of CAG- and CAA-enriched sequences require further investigation. Although our findings support a link between selective translation and alternative nutrient utilization, the extent to which uridine metabolism directly spares glutamine remains to be established. Further genetic and metabolic flux analyses will help define this relationship and determine whether similar mechanisms operate in other nutrient-stressed cancers.

In summary, our findings identify QARS spermylation as a mechanism connecting polyamine metabolism, tRNA charging, and glutamine allocation in PDAC. By favoring protein synthesis while sustaining translation of enzymes involved in alternative nutrient utilization, this modification may coordinate competing metabolic demands during nutrient stress. Our study therefore expands the view of metabolism–translation coupling by showing that translational machinery can reciprocally influence nutrient utilization.

## Data Availability

All data supporting the findings of this study are available within the paper and its Supplementary Information. Source data underlying the figures are provided as a Source Data file. Additional information and datasets generated during this study are available from the corresponding author upon reasonable request.

## Acknowledgements

This work was supported by grants from the National Natural Science Foundation of China (82230087, 82372821, 82350123, 82203228, 82372589 and 82472919), the Shanghai Municipal Education Commission–Gaofeng Clinical Medicine Grant Support (20181708), the Innovative Research Team of High-Level Local Universities in Shanghai (SHSMU-ZDCX20210802), the Shanghai Pilot Program for Basic Research–Shanghai Jiao Tong University (21TQ1400225), the 111 Project (B21024) and the Shenyang Science and Technology Plan (22-101-0-22). We thank Dr Jing-li Hou for experimental assistance.

## Author contributions

Conceptualization: L.P.H., J.L., Q.L., and Z.G.Z.; Methodology: L.P.H., X.L.Z., S.H.J., S.Z., F.H., H.Z. and G.H.S.; Formal analysis: L.P.H., X.W., J.L. and Y.M.H.; Resources: Y.M.H., Y.Y., and S.H.J.; Investigation: all authors; Supervision: L.P.H., Q.L. and Z.G.Z.; Funding acquisition: L.P.H., J.L., Y.L.Z., X.M.Y., and Z.G.Z.; Writing – original draft: L.P.H., J.L., Q.L., D.X.L., and Z.G.Z.; Writing – review and editing: all authors.

## Methods

### Cell culture

Human pancreatic cancer cell lines PANC-1 and PATU8988 were obtained from the American Type Culture Collection (ATCC). The murine PDAC cell line KPC1199 was provided by Prof. Jing Xue. Primary PDAC cell lines PDA-1 and PDA-2 were established from resected PDAC tissues. Cultures were routinely tested for mycoplasma using the Mycoplasma Detection Kit (Yeasen, 40612ES25). Cells were routinely maintained in high-glucose DMEM (Gibco, 11965092) supplemented with 10% FBS (Gibco, A5669701) at 37 °C in 5% CO. Unless otherwise indicated, nutrient-limited functional assays were performed in DMEM containing 1 mM glucose and 1% FBS. Cells were washed with PBS (Gibco, 10010023).

### Reagents and treatment conditions

Spermine (Selleck, S5524) was prepared in PBS and used at 10 μM unless otherwise indicated. Difluoromethylornithine (DFMO, Selleck, S4582), BSPM (MedChemExpress, HY-13610A), GK921 (Selleck Chemicals, cat. no. S6417), 3-methyladenine (Selleck, S2767), ruxolitinib (Selleck Chemicals, S1378), cycloheximide (Selleck Chemicals, S7418) and gemcitabine (Selleck, S1149) were prepared according to the manufacturers’ instructions. Treatment concentrations and durations are specified in the corresponding figure legends; the final solvent concentration was matched across groups. For nutrient-limitation experiments, cells were washed twice with PBS and cultured in DMEM containing 1 mM glucose, 0.5 mM glutamine and 1% dialyzed FBS for 6 h before treatment.

### Cell viability assay

PDAC cells were seeded in 96-well plates at 3,000 cells per well and allowed to adhere overnight. Cells were treated with spermine with or without the TGM2 inhibitor GK921 (Selleckchem, S6417) in DMEM containing 1 mM glucose and 1% FBS. Cell viability was assessed by measuring absorbance at 450 nm after a 1-h incubation with Cell Counting Kit-8 (CCK-8; Bimake, B34302) for 1 h. Total protein content was used for normalization. Experiments were performed in three independent replicates.

### RNA interference, QARS expression and rescue

Small interfering RNAs (siRNAs) targeting human DHPS, DOHH, and QARS, together with a non-targeting control siRNA, were synthesized by Qingke. The siRNA sequences and quantitative PCR primer sequences used for knockdown validation are provided in Supplementary Table 3. PDAC cells were seeded to reach approximately 70% confluence at the time of transfection and transfected with 20 nM siRNA using Polyplus transfection reagent (101000028) according to the manufacturer’s instructions. Cells were collected 48 h after transfection for subsequent analyses. Knockdown efficiency was confirmed at both mRNA and protein levels by quantitative PCR and immunoblotting, respectively. For rescue experiments, cells were transfected with siRNA-resistant QARS expression constructs encoding wild-type QARS. The corresponding empty vector served as the control. QARS rescue constructs were introduced 24 h after siRNA transfection, and expression levels were adjusted to approximate endogenous QARS abundance.

### Site-directed mutagenesis and QARS Q216A knock-in cells

QARS Q41A, Q86A and Q216A expression constructs were generated by site-directed mutagenesis of human QARS, and all inserts were verified by Sanger sequencing. Endogenous QARS Q216A knock-in cells were generated using CRISPR-Cas9. A guide RNA targeting QARS and a single-stranded donor oligonucleotide containing the Q216A substitution were delivered by transfection. Single-cell clones were isolated by limiting dilution. Correctly edited clones were identified by amplicon sequencing and confirmed by Sanger sequencing of both alleles. At least three independently derived clones per genotype were used, and total QARS abundance and basal growth were verified before phenotypic analyses.

### Cell-penetrating peptides

CP216 consisted of an N-terminal cell-penetrating sequence (HLYVSPWGG) fused to a competitive motif encompassing QARS Q216 (NGETADQTLSLME). The matched scrambled control contained the same cell-penetrating sequence fused to LQEMTGNLDSEAT. Peptides were synthesized by Guoping Pharmaceutical using solid-phase peptide synthesis. CP216 and the scrambled control were N-terminally acetylated and purified to >95% purity by high-performance liquid chromatography. Lyophilized peptides were dissolved in sterile PBS to generate 10 mM stock solutions, aliquoted and stored at −80 °C. For cell-based experiments, peptide stocks were diluted in culture medium and added at 10 μM for 24 h unless otherwise indicated. For in vivo studies, peptides were freshly diluted in sterile PBS and administered intraperitoneally at 10 mg kg−1 every 3 d for 4 weeks unless otherwise stated. Target engagement was evaluated by immunoblotting or immunohistochemistry using the QARS Q216-spermylation-specific antibody, with total QARS abundance assessed in parallel.

### Puromycin incorporation assays

Nascent protein synthesis was assessed by the surface sensing of translation (SUnSET) assay. Cells were incubated with puromycin dihydrochloride (Selleck, S7417) at 1 μg /ml for 30 min at 37 °C. As a translation-inhibition control, parallel cultures were pretreated with cycloheximide (Selleck, S7418) at 100 μg/ml for 15 min before the addition of puromycin; cycloheximide was maintained throughout the 30-min puromycin pulse. Cultures not exposed to puromycin served as negative controls. Cells were washed twice with ice-cold PBS, lysed immediately in RIPA buffer supplemented with protease inhibitors and analyzed by immunoblotting using an anti-puromycin antibody (Abcam, ab315887; 1:1,000). Puromycin signal was integrated across the full molecular-weight range of each lane using ImageJ (version 1.54) and normalized to total-protein staining. For in vivo labeling, mice received puromycin dihydrochloride at 20 mg/kg by intraperitoneal injection. Tumors were collected 30 min after injection, snap-frozen in liquid nitrogen and processed for immunoblotting or IF as described above.

### Ribosome run-off analysis

Ribosome run-off after acute inhibition of translation initiation was used to assess translation elongation. Cells were treated with harringtonine (Selleck, S9063) at 2 μg/ml for 0, 1, 2, 3, 4 or 5 min. At the indicated time points, cycloheximide was added to a final concentration of 100 μg ml−1 for 5 min to arrest elongating ribosomes before cell collection. The 0-min samples received an equivalent volume of vehicle followed immediately by cycloheximide. Cells were immediately washed twice with ice-cold PBS containing 100 μg ml−1 cycloheximide and lysed on ice. Cytoplasmic lysates containing equal amounts of RNA were separated on 10–50% sucrose gradients as described for polysome profiling. Absorbance profiles at 254 nm were recorded continuously, and the areas corresponding to polysomes and 80S monosomes were integrated after baseline correction using GraphPad Prism. Apparent run-off rates were estimated by fitting the normalized polysome fraction to a single-phase exponential-decay model using nonlinear least-squares regression. Three independent biological experiments were performed.

### QARS aminoacylation assay

Recombinant N-terminally His -tagged human wild-type or Q216A QARS was expressed in Escherichia coli BL21(DE3) cells and purified by Ni–NTA affinity chromatography followed by size-exclusion chromatography on a Superdex 200 Increase column. Purity exceeded 95%, as determined by Coomassie-stained SDS–PAGE. Protein concentrations were measured using a BCA assay. Where indicated, purified QARS was spermylated by incubation with recombinant TGM2 at a QARS:TGM2 molar ratio of 10:1 in the presence of 1 mM spermine, 5 mM CaCl, 1 mM DTT and 50 mM Tris-HCl, pH 7.5, for 1 h at 37 °C. Parallel control reactions omitted either TGM2 or spermine. His -tagged QARS was subsequently repurified by Ni–NTA affinity chromatography, and residual spermine and other low-molecular-weight components were removed by buffer exchange using a 30-kDa-cut-off centrifugal filter. QARS spermylation was confirmed by immunoblotting using the QARS Q216-spermylation-specific antibody. Human cytoplasmic tRNAGln was generated by in vitro transcription using T7 RNA polymerase, purified by denaturing urea–PAGE and refolded before use. Standard aminoacylation reactions contained 100 nM QARS, 5 μM tRNAGln, 50 μM total L-glutamine, 2 mM ATP, 10 mM MgCl, 30 mM KCl, 1 mM DTT and 100 mM HEPES-NaOH, pH 7.2, in a final volume of 20 μl. Reactions were initiated by addition of QARS and maintained at 37 °C. At 0, 1, 2, 4 and 6 min, aliquots were applied to Whatman 3MM filter discs pre-equilibrated with ice-cold 5% trichloroacetic acid (TCA). Filters were washed three times with ice-cold 5% TCA, followed by one wash with ethanol, air-dried and analyzed by liquid-scintillation counting. Gln-tRNA^Gln^ formation was calculated from the specific activity of glutamine after subtraction of signals from reactions lacking QARS or tRNAGln.

To determine tRNA kinetics, tRNA^Gln^ concentration was varied from 0.25 to 10 μM while ATP and l-glutamine were maintained at saturating concentrations of 2 mM and 1 mM, respectively. To assess glutamine kinetics, l-glutamine was varied from 5 to 500 μM while tRNA^Gln^ was maintained at 10 μM. Initial velocities were calculated from the linear portion of the reaction and fit to the Michaelis–Menten equation using GraphPad Prism 10.

### Untargeted metabolomics of human PDAC tissues

Fresh-frozen PDAC tissues and matched adjacent non-tumor specimens (20 pairs) were weighed on dry ice and homogenized in ice-cold 80% methanol containing isotope-labeled internal standards at 20 μl mg□¹ tissue. Extracts were incubated at −20□°C for 20□min and centrifuged at 16,000g for 15□min at 4□°C. Supernatants were dried under vacuum and reconstituted in 80% acetonitrile. Metabolites were separated on an ACQUITY UPLC BEH Amide column (2.1□×□100□mm, 1.7□μm; Waters) using a water–acetonitrile gradient containing 10□mM ammonium acetate and analyzed on a Q Exactive HF-X Orbitrap mass spectrometer (Thermo Fisher Scientific) in positive- and negative-ion modes. Intensities were normalized to tissue mass and isotope-labeled internal standards, log□-transformed and subjected to paired statistical testing with Benjamini–Hochberg correction.

### Targeted polyamine quantification

Spermine, spermidine, putrescine and ornithine were quantified by isotope-dilution LC–MS/MS. Tissue and cell samples were extracted with ice-cold 80% methanol containing the corresponding stable-isotope-labeled internal standards followed by derivatization with dansyl chloride. Analytes were separated by reverse-phase liquid chromatography and detected using a QTRAP 6500+ mass spectrometer (SCIEX) in positive-ion multiple-reaction-monitoring mode. Concentrations were determined from matrix-matched calibration curves using 1/x²-weighted linear regression and normalized to wet tissue mass or cellular protein. PanIN and PDAC tissues from KPC mice were classified by a pathologist unaware of the metabolite measurements.

### Stable-isotope tracing and glutamine allocation

For cell-based isotope tracing, cells were equilibrated in glutamine-free DMEM containing 1% dialyzed FBS and 1 mM glucose and incubated with 0.5 mM ^13^C_5_,^15^N_2_-glutamine for 1 h. Polar metabolites were recovered with ice-cold 80% methanol containing L-norvaline and analyzed by high-resolution LC–MS using hydrophilic interaction chromatography. Protein pellets were washed three times with 80% methanol, hydrolyzed in 6 M HCl at 110 °C for 24 h and analyzed by LC–MS. Because acid hydrolysis converts glutamine residues to glutamate, labeling in protein-bound glutamine/glutamate was reported as protein-bound Glx. For in vivo isotope tracing, tumor-bearing mice received ^13^C_5_,^15^N_2_-glutamine intravenously as a 0.4 g/kg bolus followed by infusion at 0.012 g/kg/ min for 60 min. Blood was collected at 0, 15, 30 and 60 min, and tumors were freeze-clamped immediately after infusion. Isotopologues were corrected for natural isotope abundance using IsoCor. To estimate relative glutamine allocation among competing biosynthetic pathways, labeled metabolite contributions were calculated based on pool-size-weighted isotopic enrichment rather than individual isotopologue abundance. The translation-associated glutamine pool was represented by protein-bound ^15^N-Glx incorporation. The TCA-cycle pool was quantified using glutamine-derived labeling of α-ketoglutarate, citrate, aspartate, succinate, fumarate and malate, with emphasis on corrected M+5 enrichment of α-ketoglutarate as a direct indicator of glutamine carbon entry. The nucleotide synthesis pool was determined from corrected isotopologue distributions of purine and pyrimidine nucleotides, including IMP, AMP, GMP, UMP and CMP. Glutamine uptake and metabolite pool sizes were evaluated separately.

### Stable isotope tracing with ^13^C_5_ uridine

To determine the metabolic fate of uridine-derived carbon, stable isotope tracing experiments were performed using uniformly labeled [U-¹³C□] uridine (Cambridge Isotope Laboratories, CLM-3680-PK). For *in vitro* tracing experiments, PDAC cells were cultured under the indicated conditions and incubated with tracer-containing medium supplemented with 0.1 mM [U-¹³C□]uridine for 30 min. Cells were rapidly washed with ice-cold PBS to remove extracellular tracer, immediately quenched with liquid nitrogen, and extracted using 80% methanol pre-chilled at −80°C. Metabolite extracts were centrifuged at 15,000 × g for 15 min at 4°C, and supernatants were collected, dried by vacuum centrifugation, and resuspended in LC–MS loading buffer for analysis. For *in vivo* tracing experiments, orthotopic PDAC-bearing KPC mice treated with scrambled CPP or CP216 were administered [U-¹³C]uridine by intraperitoneal injection (0.4 g/kg). Tumors were harvested 1 h after tracer administration, immediately freeze-clamped in liquid nitrogen, and stored at −80°C until metabolite extraction. Tumor metabolites were extracted using the same methanol-based extraction procedure described above. Extracted metabolites were analyzed using an Agilent liquid chromatography–mass spectrometry platform equipped with a high-resolution mass spectrometer (Agilent 6545 Q-TOF). Chromatographic separation was performed using Agilent 1290 Infinity II UHPLC. Metabolites were annotated based on accurate mass, retention time, and comparison with authentic standards. Raw LC–MS data were processed using Agilent MassHunter Workstation software for peak detection, integration, retention-time alignment and metabolite quantification. Isotopologue distributions were extracted based on the theoretical isotope patterns of individual metabolites.

### Colony-forming assay

PDAC cells were seeded in 35-mm dishes at 1,000 cells per dish. After 24 h, cultures were treated with spermine with or without the TGM2 inhibitor GK921 (Selleckchem, S6417) at the indicated concentrations for 2 weeks in DMEM containing 1 mM glucose and 1% FBS. Colonies were fixed in 4% paraformaldehyde and stained with 0.05% crystal violet. Colonies were counted using ImageJ.

### 3D on-top growth

Three-dimensional on-top cultures were established by polymerizing the extracellular matrix (ECM) before seeding cells. For ECM preparation, growth factor-reduced Matrigel (Corning) was thawed at 4 °C overnight and mixed thoroughly with ice-cold serum-free DMEM at a 1:1 ratio (final protein concentration, 4–6 mg ml−1). This mixture was immediately dispensed into pre-cooled 24-well plates (100 μl per well) and polymerized at 37 °C for 30–45 min in a humidified CO□incubator. Tumor cells were suspended in low-glucose medium containing 1% FBS at a density of 1 × 10□cells ml−1. Cell suspensions (500 μl per well) were carefully pipetted onto the polymerized matrix surface to avoid structural disruption. Cells were allowed to attach without agitation for 15–30 min at 37 °C before the addition of 1 ml fresh medium per well. Cultures were maintained under the indicated treatment conditions for 14 d after seeding. Culture morphology was documented periodically by phase-contrast imaging using an inverted microscope (Nikon Eclipse Ti) equipped with a 20×/0.40 NA objective.

### Organoid culture and assay

Surgically resected primary PDAC tissues were minced into 1–2 mm³ fragments using sterile scalpels. Fragments were digested in Advanced DMEM/F12 (Gibco) supplemented with 10% fetal bovine serum (FBS; Gemini Bio), 2 mM L-glutamine (Corning), 10 μM Y-27632 (ROCK inhibitor, Selleckchem), 500 nM A83-01 (TGF-β inhibitor, Tocris), 100 μg ml−1 primocin (InvivoGen) and 5 mg ml−1 collagenase II (Worthington). The digestion mixture was incubated at 37 °C for 90–120 min with gentle agitation (50 rpm) in a cell-culture incubator (5% CO□, 95% humidity). After digestion, cell suspensions were passed through 100-μm cell strainers (Falcon) and centrifuged at 300 × g for 5 min at 4 °C. Pelleted cells were resuspended in ice-cold Cultrex UltiMatrix Reduced Growth Factor Basement Membrane Extract (BME; R&D Systems, BME001-05) at a density of 5–10 × 10³ viable cells μl−1, as determined by trypan blue exclusion. BME–cell suspensions (30-μl droplets) were plated in pre-warmed 24-well plates (Corning) and polymerized at 37 °C for 30 min before overlay with 1 ml PancreaCult Organoid Growth Medium (STEMCELL Technologies, 100-0820) containing 1× penicillin–streptomycin (Gibco). Organoids were routinely passaged every 7–10 d using Gentle Cell Dissociation Reagent (STEMCELL Technologies). For treatment assays, organoids were cultured in medium containing 1 mM glucose and 1% FBS and treated with spermine with or without the TGM2 inhibitor GK921 (Selleckchem, S6417), as indicated. Short tandem repeat (STR) profiling showed ≥85% allele matching across 16 STR loci.

### Western blotting

Cells or tumor tissues were lysed on ice using WB buffer (NCM Biotech, P70100) supplemented with protease/phosphatase inhibitor cocktail. Lysates were incubated on ice for 10 min and centrifuged (18,000 × g, 10 min, 4 °C) to collect the supernatants. Protein concentrations were measured using the Pierce BCA Assay (Thermo Fisher, 23227). Equal protein amounts (20–25 μg), together with NcmColor ladder (NCM Biotech, P9001), were resolved on 4–12% Bis-Tris gels (Thermo Fisher, NP0336BOX) at 120 V. Proteins were transferred onto methanol-activated PVDF membranes (Millipore) at 25 V for 1 h. Membranes were blocked with 5% non-fat milk in TBST (0.1% Tween-20) for 1 h and then incubated overnight at 4 °C with primary antibodies against QARS (ZENbio, 671631; 1:1,000), spermylated QARS Q216 (generated in this study; 1:1,000), UPP1 (Proteintech, 14186-1-AP; 1:1,000) and β-actin (Cell Signaling Technology, 4970; 1:3,000). After three 10-min washes with TBST, membranes were incubated with HRP-conjugated secondary antibodies (Cell Signaling Technology, 7074S, anti-rabbit; 7076P2, anti-mouse; 1:10,000) for 1 h. Chemiluminescence was detected using Clarity Max substrate (Bio-Rad, 705062), and images were acquired on a ChemiDoc system (Bio-Rad) using Image Lab 2.4.

### Quantitative tRNA aminoacylation assay

tRNA aminoacylation was measured using a modified version of a previously described high-sensitivity assay. PDAC cells treated with spermine or vehicle were lysed using TriPure RNA Isolation Reagent (Roche, 11667157001), followed by overnight precipitation with ice-cold absolute ethanol (1:1, v/v) at −80 °C. RNA pellets were washed with 70% ethanol and resuspended in 0.3 M sodium acetate buffer (pH 4.5) containing 10 mM EDTA, then reprecipitated by ethanol-gradient purification. Total RNA (2 μg per sample) was divided into two groups. For the oxidation group, RNA was treated with 10 mM sodium periodate (NaIO ; Sigma-Aldrich) in the dark at 25 °C for 20 min. Control samples were processed with equimolar NaCl. Reactions were quenched with 100 mM D-glucose (Sigma-Aldrich). Samples were supplemented with 5 pmol Glu-tRNA internal control (Sigma-Aldrich, R4018) before secondary precipitation with isopropanol. The final RNA pellets were resuspended in diethyl pyrocarbonate (DEPC)-treated water.

### Northern blot

The aminoacylation status of Gln-tRNA was assessed by acid–urea polyacrylamide gel electrophoresis (acid–urea PAGE) followed by Northern blotting using the Biotin Northern Blot Kit (for Small RNA) (Beyotime Biotechnology, R0220). Total RNA was isolated from PDAC cells using TRIzol reagent (Invitrogen, 15596026) under acidic conditions to preserve aminoacyl-tRNA ester bonds. RNA samples were maintained at 4°C and resuspended in sodium acetate buffer (pH 5.0). Equal amounts of RNA (1.5 μg per lane) were mixed with acid–urea loading buffer and separated on 6.5%polyacrylamide gels containing 8 M urea and 100 mM sodium acetate (pH 5.0). Electrophoresis was performed at 4°C using sodium acetate running buffer (pH 5.0) under 100 V for 1 h to resolve aminoacylated and deacylated Gln-tRNA. RNA was transferred to positively charged nylon membranes (Hybond-N+, GE Healthcare, RPN203B) in 0.5× TBE buffer at 0.2 A for 60 min using a Trans-Blot SD Semi-Dry Transfer Cell (Bio-Rad, 1703940). UV crosslinking was performed at a dose of 1,200 × 100 μJ cm−2 using a Spectrolinker XL-1000 UV Crosslinker (Invitrogen, C300017). Membranes were hybridized with a biotin-labeled oligonucleotide probe complementary to tRNA^Gln^ using BeyoHyb Quick Hybridization Solution according to the manufacturer’s instructions. After sequential low- and high-stringency washes, hybridized probes were detected using streptavidin–horseradish peroxidase and Blots were exposed to Amersham Hyperfilm ECL (GE Healthcare, 28906839) for 5–30 min. Signals were acquired using a ChemiDoc imaging system (Bio-Rad), and band intensities were quantified using ImageJ. The Gln-tRNA charging fraction was calculated as the intensity of the aminoacylated Gln-tRNA band divided by the combined intensities of aminoacylated and deacylated Gln-tRNA bands.

### Histology, immunohistochemistry and scoring

Formalin-fixed, paraffin-embedded human and mouse PDAC tissues were sectioned at 4□μm using a rotary microtome (Leica RM2255). Sections were deparaffinized in xylene, rehydrated through graded ethanol and subjected to antigen retrieval in 10□mM sodium citrate buffer (pH 6.0) at 95□°C for 20□min. Endogenous peroxidase activity was quenched with 3% H□O□for 10□min, and sections were blocked with 5% BSA in TBST for 1 h. Sections were incubated overnight at 4 °C with primary antibodies against QARS Q216 spermylation (custom rabbit antibody), total QARS/GlnRS (Proteintech, 12645-1-AP), ODC1 (Proteintech, 17003-1-AP), AMD1 (Proteintech, 11052-1-AP), SMS (Proteintech, 15979-1-AP), PAOX (Proteintech, 18972-1-AP), UPP1 (Proteintech, 14186-1-AP), UCK2 (Proteintech, 10511-1-AP), MYC (Cell Signaling Technology, 5605), PCNA (Cell Signaling Technology, 13110), human Ki-67 (Cell Signaling Technology, 9449) or mouse Ki-67 (Cell Signaling Technology, 12202), each diluted 1:100. Signal was developed using the appropriate SignalStain Boost HRP polymer and DAB substrate (Cell Signaling Technology), followed by hematoxylin counterstaining. H&E staining was performed using standard procedures. Whole-slide images were acquired using an Aperio AT2 scanner and analyzed in QuPath (version 0.5.1). PanIN and PDAC regions were annotated by a pathologist blinded to genotype and treatment, and the positive area or positive-cell fraction was quantified using a fixed threshold in five non-overlapping fields per sample. For clinical specimens, staining was scored independently by two pathologists blinded to clinical outcomes. H-scores were calculated as the sum of staining intensity (0–3) multiplied by the percentage of positive tumor cells, yielding scores of 0–300; discrepancies exceeding 30 points were resolved by consensus.

### Immunofluorescence

Antigen retrieval was performed on PDAC tissue sections in 10 mM sodium citrate buffer (pH 6.0; Sigma, C9999) at 95 °C for 20 min using a PT Module system (Thermo Fisher). Samples were permeabilized with 0.5% Triton X-100 for 15 min at room temperature and then blocked for 1 h in PBS containing 0.2% Triton X-100, 0.05% Tween-20, 1% BSA (Sigma, A7906) and 5% normal donkey serum (Jackson ImmunoResearch, 017-000-121). Primary antibodies diluted in antibody stabiliser (Dako, S3022) were applied overnight at 4 °C in humidified chambers: QARS (Proteintech, 12645-1-AP; 1:200), spermine antibody (Abcam, ab26975; 1:100), spermylated QARS Q216 (generated in this study; 1:100) and UPP1 (Proteintech, 14186-1-AP; 1:100). After three washes with PBS-T (0.1% Tween-20), sections were incubated with species-matched secondary antibodies conjugated to Alexa Fluor dyes (Invitrogen; 1:1,000) for 2 h protected from light. Samples were mounted with ProLong Diamond Antifade (Invitrogen, P36965) and imaged on a Zeiss LSM 880 confocal microscope. Images were quantified using Imaris 9.7 (Bitplane) with automated spot detection (minimum diameter, 0.3 μm; quality threshold >50). Co-localization was assessed using Pearson’s correlation coefficient in the ZEN Coloc module, with a threshold of >0.6.

### Identification of spermylated proteins

An alkyne-functionalized spermine analog was synthesized and characterized by LC–MS and NMR. Cells were incubated with 10 μM alkyne-functionalized spermine for 24 h, washed with ice-cold PBS and lysed. Probe-labeled proteins were biotinylated to azide–PEG4–biotin by copper(I)-catalyzed azide–alkyne cycloaddition using 100 μM azide–PEG4–biotin, 1 mM CuSO□, 1 mM THPTA and 5 mM sodium ascorbate for 1 h at room temperature. Proteins were precipitated to remove excess click reagents, resolubilized and incubated with streptavidin magnetic beads overnight at 4 °C. After stringent washing, enriched proteins were digested on-bead with trypsin and analyzed by LC–MS/MS. Probe-free and excess-spermine competition samples were included as negative controls.

### Glutamine-codon reporters and synonymous recoding

Firefly luciferase reporters containing 0, 6, 12, or 18 glutamine codons were constructed by inserting synthetic coding sequences immediately upstream of the stop codon in a pcDNA3.1 firefly luciferase vector. CAA- and CAG-enriched reporter pairs encoded identical amino-acid sequences and differed only in the synonymous glutamine codons. The coding sequences of UPP1, UCK2 and MYC were synonymously recoded to encode glutamine residues predominantly with either CAA or CAG. Overall GC content and codon adaptation index were maintained within 2% and 5%, respectively, between paired constructs, without altering the amino-acid sequence, untranslated regions or epitope tags. Cells were co-transfected with firefly reporter and pRL-TK Renilla luciferase plasmids at a 10:1 ratio using Lipofectamine 3000 (Thermo Fisher Scientific). At 24 h after transfection, cells were treated with spermine, CP216, scrambled peptide or vehicle at 10 μM for a further 24 h. Luciferase activity was measured using the Dual-Luciferase Reporter Assay System (Promega, E1910) on a GloMax luminometer. Firefly luciferase activity was normalized to Renilla activity. Expression of the recoded UPP1, UCK2 and MYC constructs was evaluated by immunoblotting, whereas transcript abundance was measured by RT–qPCR to distinguish translational regulation from changes in mRNA abundance.

### Ribosome sucrose density gradient sedimentation

PDAC cells were cultured under the indicated conditions and harvested at approximately 80% confluence. Cycloheximide (CHX, 100 μg ml−1) was added to the culture medium for 15 min at 37 °C to arrest ribosomes on actively translating mRNA. Cultures were washed with ice-cold PBS containing 100 μg ml−1 CHX and lysed in hypotonic buffer (10 mM HEPES, pH 7.4, 1.5 mM MgCl□, 10 mM KCl, 1 mM DTT, 0.5% Triton X-100 and protease inhibitors). Lysates were homogenized using a Dounce homogenizer and clarified by centrifugation at 720 × g for 5 min to remove nuclei and debris. A discontinuous sucrose gradient was prepared in ultracentrifuge tubes by layering sucrose solutions (10%, 20%, 30%, 40% and 50%, w/v, in PBS) using a gradient mixer. The gradient was stabilized at 4 °C for 30 min to form sharp interfaces. Clarified lysates (equivalent to 1–2 × 10□cells) were layered onto the gradient without disturbing the sucrose layers. Tubes were centrifuged at 100,000 × g for 16 h at 4 °C using an SW 40 Ti rotor (Beckman Coulter). Ribosomal complexes (monosomes and polysomes) were thereby separated on the basis of buoyant density. Gradients were fractionated into 300-μl aliquots from top to bottom using a peristaltic pump with continuous monitoring at 260 nm. Ethanol was added to each fraction, and samples were mixed and stored at −20 °C overnight. Fractions were centrifuged at 18,000 × g for 15 min at 4 °C, and pellets were dissolved in 1× SDS–PAGE sample buffer and heated at 99 °C for 3 min.

### Ribosome profiling (Ribo-seq)

Primary human PDAC cells were immediately washed with ice-cold PBS containing cycloheximide (100 μg ml−1) and lysed in ribosome-profiling buffer containing 20 mM Tris-HCl (pH 7.4), 150 mM NaCl, 5 mM MgCl□, 1 mM dithiothreitol, 1% Igepal CA-630, 100 μg ml−1 cycloheximide and RNase inhibitor. Lysates were clarified by centrifugation at 20,000 × g for 10 min at 4 °C, and equal amounts of RNA were used for ribosome-footprint preparation. Clarified lysates were digested with 6.25 U RNase I (Epicentre) in a final volume of 50 μl containing 0.1% Igepal CA-630 for 45 min at room temperature. Digestion was terminated with RNase inhibitor, and samples were layered onto 50 μl of 1 M sucrose cushion prepared in ribosome-profiling buffer. Ribosomes were pelleted by ultracentrifugation at 165,000 × g for 30 min at 4 °C using a TLA-100 rotor (Beckman Coulter). RNA was extracted from ribosome pellets using 700 μl QIAzol, followed by chloroform extraction and ethanol precipitation. Ribosome-protected fragments of 28–35 nucleotides were isolated on 15% TBE–urea polyacrylamide gels. Gel slices were crushed and incubated in 400 μl RNA extraction buffer containing 300 mM sodium acetate (pH 5.5), 1 mM EDTA and 0.25% SDS. Eluates were passed through Spin-X filters (Corning, 8160), and RNA was precipitated with 2.5 volumes of ethanol and 1.5 μl GlycoBlue (Thermo Fisher Scientific, AM9516).

Ribosome-profiling libraries were prepared using the D-Plex Small RNA-seq Kit (Diagenode) according to the manufacturer’s instructions. Libraries were amplified for 10 PCR cycles, purified using a PCR purification kit (Qiagen) and size-selected on a 3% agarose cassette using the BluePippin system (Sage Science), with a collection range of 180–235 bp and a target size of 208 bp. Library fragment sizes and concentrations were assessed using the Agilent High Sensitivity DNA Kit. Libraries were pooled at equimolar concentrations and sequenced on an Illumina NovaSeq 6000 SP flow cell in paired-end mode.

Ribosome-profiling reads were processed with RiboFlow. The first 12 nucleotides of read 1 were extracted as unique molecular identifiers, and the following four template-switching nucleotides matching NGGG were removed. The 3′ adapter sequence (AAAAAAAAAACAAAAAAAAAA) and low-quality bases were trimmed, and PCR duplicates were collapsed using the unique molecular identifiers. Reads shorter than 20 nt after trimming were discarded. Reads mapping to human rRNA, tRNA, mitochondrial rRNA or other abundant non-coding RNAs were removed. Remaining reads were aligned to the human GRCh38 reference genome and GENCODE human gene annotation using STAR. MANE Select transcripts were used where available; for genes without a MANE Select annotation, the protein-coding transcript with the longest coding sequence was retained. Transcripts were not selected randomly. P-site offsets were determined separately for each footprint length using metagene periodicity around annotated translation start sites. Only footprints showing the expected coding-region enrichment and three-nucleotide periodicity were retained for downstream analyses. Gene-level ribosome occupancy was calculated from P-site counts within coding sequences and normalized as reads per million mapped footprints. The ribosome-profiling data were deposited in the Sequence Read Archive under BioProject accession PRJNA1261636.

Matched RNA-seq libraries were generated from the same biological samples used for ribosome profiling. After adapter and quality trimming, RNA-seq reads were aligned to the human GRCh38 reference genome with STAR using the same GENCODE gene annotation, and gene-level mRNA abundance was quantified from uniquely mapped reads. Genes with low read counts in either assay were excluded before differential analysis. Translational efficiency (TE) was defined as ribosome-footprint abundance relative to the matched mRNA abundance. Differential TE was evaluated using a generalized linear model containing treatment, assay type (Ribo-seq versus RNA-seq) and their interaction; the assay-by-treatment interaction represented the treatment-dependent change in TE. Multiple-testing correction was performed using the Benjamini-Hochberg method, with a false-discovery rate (FDR) of <0.05 considered significant. For codon-resolved analyses, A-site positions were assigned to individual coding codons after footprint-length-specific offset correction. Codon pause scores were calculated as A-site occupancy at a given codon normalized to the mean A-site density across the coding sequence of the same transcript. For the CAG-density analysis, CAG frequency was expressed as the number of CAG codons per 100 coding codons within each analyzed coding sequence.

### Real-time PCR

Total RNA was isolated according to the manufacturer’s instructions, treated with DNase I (RNase-free) and reverse-transcribed into cDNA with random hexamers using the SuperScript II First-Strand Synthesis System (Invitrogen, 18064071). qPCR reactions were performed in technical triplicate using an Applied Biosystems ViiA 7 Real-Time PCR system as previously described. Reactions contained cDNA generated by reverse transcription of 25 ng total RNA, 150 nM of each primer and 5 μl 2× JumpStart SYBR Green PCR mix (Invitrogen) in a total volume of 10 μl. Relative mRNA levels were calculated using the comparative Ct (cycle-threshold) method and normalized to cyclophilin.

### tRNA qPCR

Total RNA was extracted using the TriPure isolation method (Roche, 11667157001). At least 3 μg total RNA was used for RNA demethylation, followed by RNA reprecipitation and first-strand cDNA synthesis using the rtStar tRNA Pretreatment and First-Strand cDNA Synthesis Kit (Arraystar, AS-FS-004), according to the manufacturer’s instructions. Individual tRNAs were quantified by qPCR using SYBR Green and pre-designed tRNA primer sets (Arraystar) on a LightCycler 480 System (Roche). qPCR was performed with an initial denaturation at 95 °C for 10 min, followed by 40 cycles of 95 °C for 10 s and 60 °C for 1 min, and a final melting-curve analysis.

### Measurement of aminoacyl-tRNA levels using the PARTI assay

The abundance of aminoacylated Gln-tRNA was quantified using the periodate-based tRNA charging assay (PARTI) with modifications according to the published protocol. Briefly, total RNA was isolated from PDAC cells under conditions preserving aminoacyl-tRNA ester bonds. Purified RNA was subjected to sodium periodate (NaIO) oxidation, which selectively modifies the 3′-terminal ribose of uncharged tRNAs, whereas aminoacylated tRNAs are protected from oxidation by the attached amino acid.

Following periodate oxidation and quenching, RNA samples were enzymatically digested to release amino acids and nucleotide-derived products. The resulting derivatives were analyzed by liquid chromatography–tandem mass spectrometry (LC–MS/MS) to quantify aminoacyl-tRNA-derived amino acid signatures. The abundance of charged Gln-tRNA was determined based on the enrichment of glutamine-derived signals from periodate-resistant tRNA molecules. Untreated RNA samples and deacylated controls were analyzed in parallel to define total tRNA abundance and background signals, respectively.The relative charging level of Gln-tRNA was calculated as the ratio of aminoacylated tRNA-derived glutamine signal to total tRNA-associated signal after normalization to internal standards.

### Generation of QARS Q216^spm^ antibody

An antibody recognizing QARS spermylated at Q216 was generated in collaboration with PTM BIO. Two Q216-spermylated peptides (NGETAD-Q(Spermine)-TLSLME-C and C-NGETAD-Q(Spermine)-TLSL) and an unmodified control peptide (C-NGETAD-Q-TLSLME) were synthesized and verified by LC–MS/MS. The modified peptides were conjugated to keyhole limpet hemocyanin (KLH). Three New Zealand rabbits were immunized with each modified peptide on days 0, 21, 35 and 54. Serum was collected on days 45, 50, 64 and 69, antibody titers were assessed by ELISA and immunoblotting, and the optimal serum was selected for antibody purification. Antibody specificity was evaluated by peptide dot blot and immunoblotting. For peptide dot blots, Q216-spermylated and unmodified QARS peptides were spotted in a dilution series onto nitrocellulose membranes. After drying for 30 min, membranes were blocked with 5% BSA, incubated with primary antibody (1:2,000) followed by secondary antibody (1:10,000) for 1 h each, and visualized using an enhanced chemiluminescence (ECL) kit (Abcam). For immunoblot validation, spermine (5 mmol l−1) was transamidated to QARS protein (Novus; 10 μg) in the presence of 0.25 μg functional TGM2 protein (Novus) in a final volume of 25 μl enzymatic buffer containing 25 mM Tris-Cl (pH 8.0), 5 mM CaCl□and protease inhibitors. Protein samples were separated by 8–10% SDS–PAGE and transferred to 0.22-μm nitrocellulose membranes. After blocking with 5% BSA in TBST for 1 h at room temperature, membranes were incubated overnight with primary antibody (1:1,000) and for 2 h with secondary antibody (Immunoway; 1:10,000). Blots were visualized using the Bio-Rad imaging system.

### Enzymatic spermylation assays

MDC (Sigma, 30432; 5 μM) or spermine (5 μM) was transamidated to QARS protein (Novus, NBP2-52622; 10 μg) by 0.25 μg functional TGM2 protein (Novus, 5418-TG-010) in a final volume of 25 μl enzymatic buffer (25 mM Tris-Cl, pH 8.0, 5 mM CaCl□and protease inhibitors). For MDC assays, reactions were treated with the TGM2 inhibitor GK921 (selleck; 4 mM) or competed with excess spermine (500 μM). Reactions were incubated at 37 °C for 3 h in the dark and then heated to 98 °C for 8 min in denaturing sample buffer before gel electrophoresis. Spermylated proteins were visualized under UV light, followed by Coomassie staining to assess protein loading. QARS protein was synthesized by Jiangsu JiTai Peptide Industry Science and Technology Co., Ltd.

### Spermylation LC-MS/MS analyses

For LC–MS/MS, proteins were separated by 10% SDS–PAGE and stained with Coomassie Brilliant Blue. Selected protein bands were excised and digested with trypsin (Promega) for 20 h at 37 °C. Peptides were desalted using C18 StageTips and separated on an Easy-nLC 1200 system (Thermo Fisher) at a flow rate of 300 nl min−1. Mobile phase A was 0.1% formic acid in water; mobile phase B was 0.1% formic acid in 95% acetonitrile. The column was equilibrated with 100% mobile phase A. The gradient was 2–8% B from 0–2 min, 8–28% B from 2–42 min, 28–40% B from 42–50 min, 40–100% B from 50–51 min and 100% B from 51–60 min. Peptides were analyzed by data-dependent acquisition on a Q Exactive HF-X mass spectrometer (Thermo Fisher) in positive-ion mode over a precursor m/z range of 300–1,500. Full MS scans were acquired at a resolution of 60,000 at m/z 200, with an AGC target of 3 × 10 and a maximum injection time of 50 ms. The 20 most intense precursor ions from each full scan were selected for MS/MS. MS2 spectra were acquired at a resolution of 15,000 at m/z 200, with an AGC target of 1 × 10□, a maximum injection time of 50 ms, HCD activation and an isolation window of 1.6 m/z. Database searching was performed using MaxQuant 1.6.1.0 against the UniProt human proteome database (Homo sapiens).

### Human PDAC samples

Human PDAC samples for immunohistochemical staining were obtained from the Department of Biliary-Pancreatic Surgery, Ren Ji Hospital, School of Medicine, Shanghai Jiao Tong University. Clinical data included age, sex, diagnosis, tumor location, disease stage and histological differentiation. All procedures involving human samples were approved by the Ethics Committee of Ren Ji Hospital, School of Medicine, Shanghai Jiao Tong University (KY2023-111-C), and written informed consent was obtained from all participants.

### Animal studies

All animal procedures were approved by the Institutional Animal Care and Use Committee of Shanghai Jiao Tong University (IACUC A2023083). For CP216 pharmacokinetic and tolerability assessments, six-week-old male C57BL/6J mice received CP216 (10 mg kg−1, intraperitoneally) in sterile PBS. For pharmacokinetic analysis, blood was collected from three mice per time point before dosing and at 0.25, 0.5, 1, 2, 4 and 8 h after administration. Plasma CP216 was quantified by LC–MS/MS following acetonitrile precipitation, and pharmacokinetic parameters were calculated by non-compartmental analysis. For tolerability assessment, mice received CP216 or scrambled control peptide (10 mg kg−1, intraperitoneally, every 3 d for 4 weeks; n = 5 per group). Body weight was recorded every 2–3 d. At the endpoint, serum alanine aminotransferase, aspartate aminotransferase, blood urea nitrogen and creatinine were measured using an automated clinical chemistry analyzer. Liver, kidney, heart, lung, spleen and colon were collected, fixed in 4% neutral-buffered formalin and subjected to H&E staining. Histopathology was evaluated by a pathologist blinded to treatment allocation.

To generate subcutaneous xenografts, PDA-1 cells were suspended in a 1:1 mixture of serum-free DMEM and Matrigel (Corning, 354234), and 1 × 10 cells in 100 μl were injected into the flank. Once tumors reached approximately 50 mm³, mice were stratified by tumor size and randomly assigned to treatment. Patient-derived xenografts were established from de-identified, pathology-confirmed PDAC specimens. Tumor fragments of approximately 2 mm³ were implanted subcutaneously into 6-week-old male NOD-SCID mice. When PDX tumors reached approximately 80 mm³, mice were randomly assigned to CP216 or scrambled control peptide (10 mg kg−1, intraperitoneally, every 3 d) for 4 weeks. Tumors were measured every 3 d with callipers, and volume was calculated as V = L × W²/2. Investigators measuring tumors were blinded to treatment allocation. Mice were euthanized when tumors reached 1,500 mm³, became ulcerated or interfered with normal activity, or when animals exhibited >20% body-weight loss or persistent signs of distress. No animals were excluded unless they met prospectively defined humane or technical exclusion criteria.

For orthotopic tumor implantation, mice were anesthetized with inhaled isoflurane, the left flank was prepared aseptically, and 50,000–100,000 cells in 25 μl were injected into the pancreas before wound closure. Orthotopic tumors were monitored weekly by bioluminescence imaging. Mice received D-luciferin potassium salt (ShareBio, SB-D1009; 150 mg kg−1, intraperitoneally) and were imaged 10 min later under 1.5–2% isoflurane anesthesia using an IVIS Spectrum system. Total photon flux was quantified within identically sized abdominal regions of interest using Living Image software. Orthotopic experiments were generally concluded 3–4 weeks after implantation. LSL-*Kras*^G12D/+^, LSL-*Trp53*^R172H/+^ and *Pdx1*-Cre mice were obtained from The Jackson Laboratory to generate KPC mice. For therapeutic studies, KPC mice were randomly assigned to vehicle, CP216 (10 mg kg−1), gemcitabine (50 mg kg−1) or CP216 plus gemcitabine. CP216 and gemcitabine were administered intraperitoneally in PBS every 3 d. MRI was performed using a 7-T Bruker BioSpec small-animal system equipped with a 40-mm mouse body volume coil. Mice were maintained under 1.5–2% isoflurane anesthesia, with respiratory rate and body temperature monitored throughout imaging. Tumors were manually segmented on T2-weighted images using ITK-SNAP (version 3.8.0) by an observer blinded to treatment allocation. Tumor volume was calculated as the summed tumor area across sections multiplied by slice thickness.

To generate QARS Q216A-edited pancreatic tumors in vivo, an AAV-mediated CRISPR editing strategy was used to introduce the Q216A point mutation into the endogenous Qars locus. The editing design was based on homology-directed repair to convert the endogenous glutamine codon encoding Q216 into an alanine codon while preserving endogenous QARS expression.The selected sgRNA sequence GTCTCTGATGGAGCAGCTCC-GGG was cloned into an AAV-compatible CRISPR vector containing SpCas9 nuclease. For HDR-mediated editing, AAV HDR donor template was designed containing the Qars Q216A substitution (CAG to GCG) flanked by left and right homology arms derived from the endogenous Qars locus. Recombinant AAV8 vectors carrying the CRISPR editing components and HDR donor template were generated by Laiying gene technology company shanghai, were administered systemically to KPC mice to achieve pancreatic tumor editing. Following tumor establishment, mice were monitored until experimental endpoints. Tumors were harvested for genomic DNA extraction, molecular validation and phenotypic analysis.

Editing efficiency was assessed by PCR amplification of the targeted Qars genomic region followed by Sanger sequencing. Mice were housed in a specific-pathogen-free facility (maximum five per cage) under a 12-h light/12-h dark cycle, 30–70% humidity and 20–23 °C, with chow and water available ad libitum.

### Statistical analysis

Quantitative data are presented as the mean ± standard deviation (SD). Comparisons between two groups used two-sided paired t-tests for matched tissues and two-sided unpaired t-tests for independent samples when model assumptions were met. Comparisons among more than two groups used one- or two-way ANOVA with the stated multiplicity correction. Longitudinal tumor-growth and body-weight data were analyzed using a mixed-effects model with mouse as a random effect and time, treatment and their interaction as fixed effects. Survival outcomes were analyzed by the Kaplan–Meier method and a two-sided log-rank test; hazard ratios and 95% confidence intervals were estimated using Cox proportional-hazards models. Categorical clinicopathological variables were analyzed by χ² or Fisher’s exact test, as appropriate. Correlations were assessed using Pearson’s correlation coefficient. Normality was assessed using the Shapiro–Wilk test and Q–Q plots, and homogeneity of variance was evaluated using the Brown–Forsythe test. Exact P values, test statistics, degrees of freedom, effect sizes and confidence intervals are reported as appropriate. Multiple-testing adjustment used the Benjamini–Hochberg method. Analyses were performed in GraphPad Prism 10. All experiments included ≥3 independent biological replicates, with exact replication numbers specified in the corresponding figure legends. A threshold of P < 0.05 defined statistical significance.

