## supplement figures for "QARS spermylation directs glutamine allocation to selective translation in cancer"

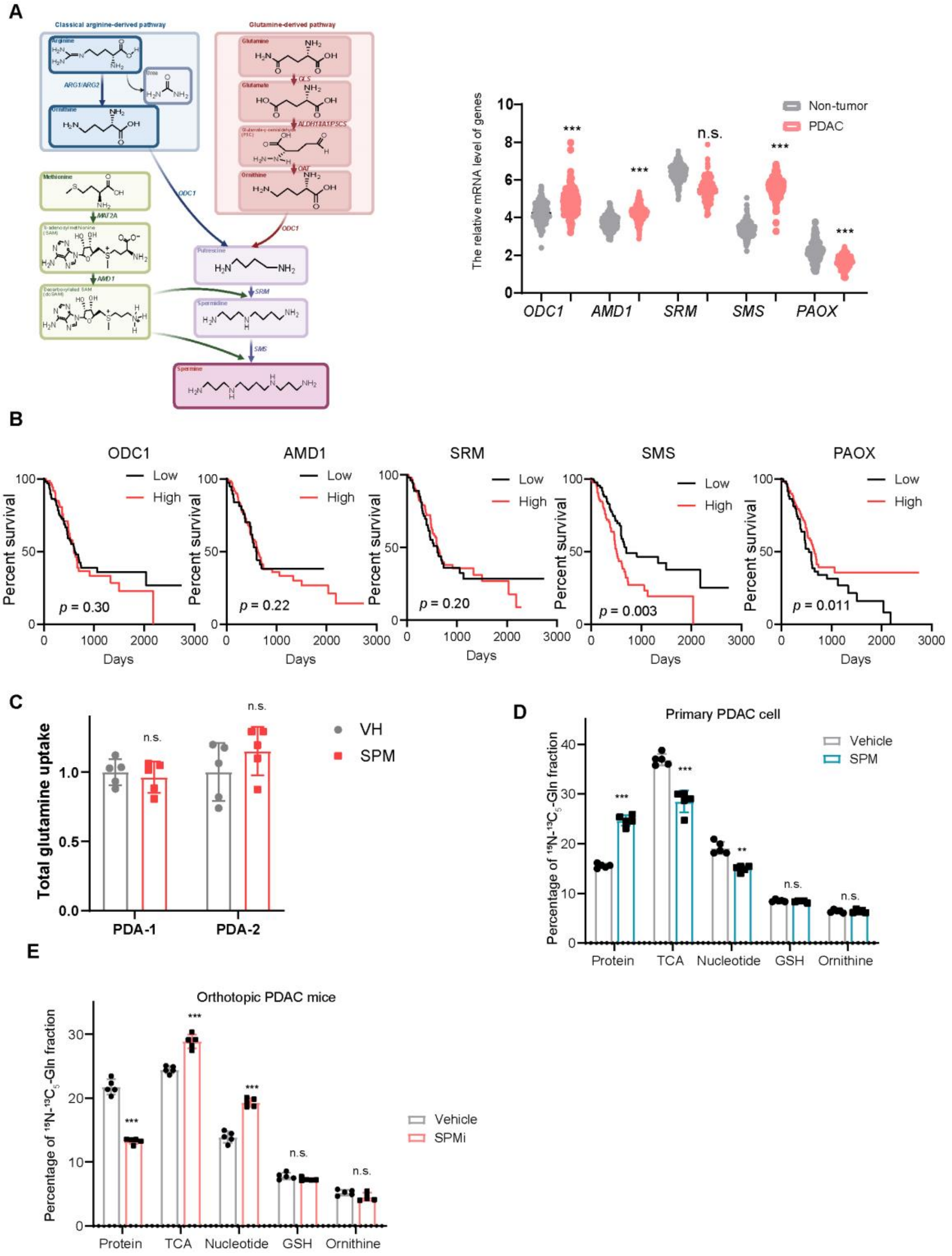


Supplement figure 1

**Spermine redistributes glutamine utilization without increasing glutamine uptake. a,** Relative expression of ODC1, AMD1, SRM, SMS and PAOX in human PDAC and non-tumour tissues. **b,** Kaplan–Meier overall-survival analyses stratified by low versus high expression of the indicated polyamine-metabolic enzymes. **c,** Total glutamine uptake in PDA-1 and PDA-2 primary PDAC cells treated with vehicle or spermine (SPM). **d,** Distribution of U-13C5,15N2-glutamine-derived label among protein, TCA-cycle intermediates, nucleotide pools, glutathione (GSH) and ornithine in primary PDAC cells treated with vehicle or SPM under nutrient-limited conditions. **e,** Corresponding glutamine-allocation analysis in orthotopic PDAC tumours in vivo. Data are mean ± s.d.; individual points represent biologically independent samples. Statistical analyses are described in Methods.

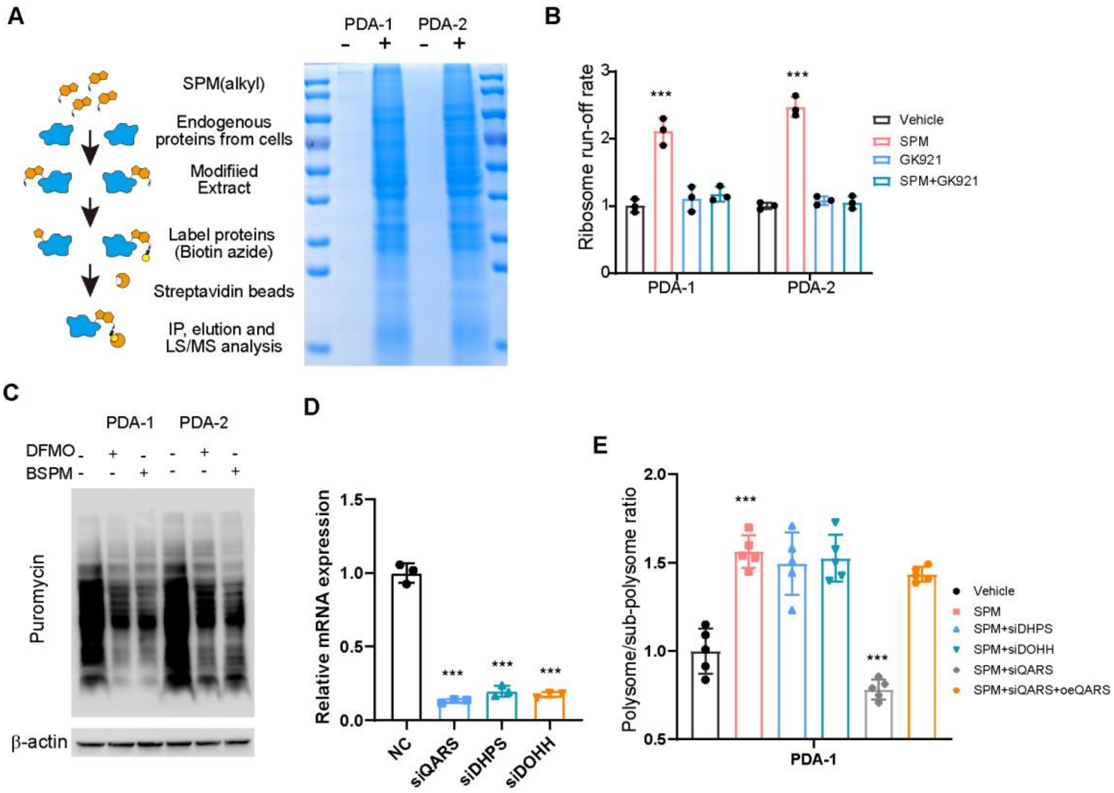


Supplement figure 2

**Chemoproteomic identification and functional validation of spermylation-dependent translational control. a,** Schematic of the alkyne-spermine chemoproteomic workflow used to enrich spermylated proteins by click conjugation to biotin and streptavidin capture, followed by LC–MS/MS; a representative protein-staining gel of labelled extracts from PDA-1 and PDA-2 cells is shown. **b,** Quantification of ribosome run-off rates in PDA-1 and PDA-2 cells treated with vehicle, SPM, the TGM2 inhibitor GK921, or SPM plus GK921. **c,** Puromycin-incorporation immunoblotting in PDA-1 and PDA-2 cells following inhibition of spermine synthesis with DFMO or BSPM; β-actin is shown as a loading control. **d,** RT–qPCR validation of siRNA-mediated depletion of QARS, DHPS and DOHH relative to the non-targeting control (NC). **e,** Polysome-to-subpolysome ratio in spermine-treated PDA-1 cells after depletion of DHPS, DOHH or QARS, with QARS re-expression in QARS-depleted cells where indicated. Data are mean ± s.d.; statistical analyses are described in Methods.

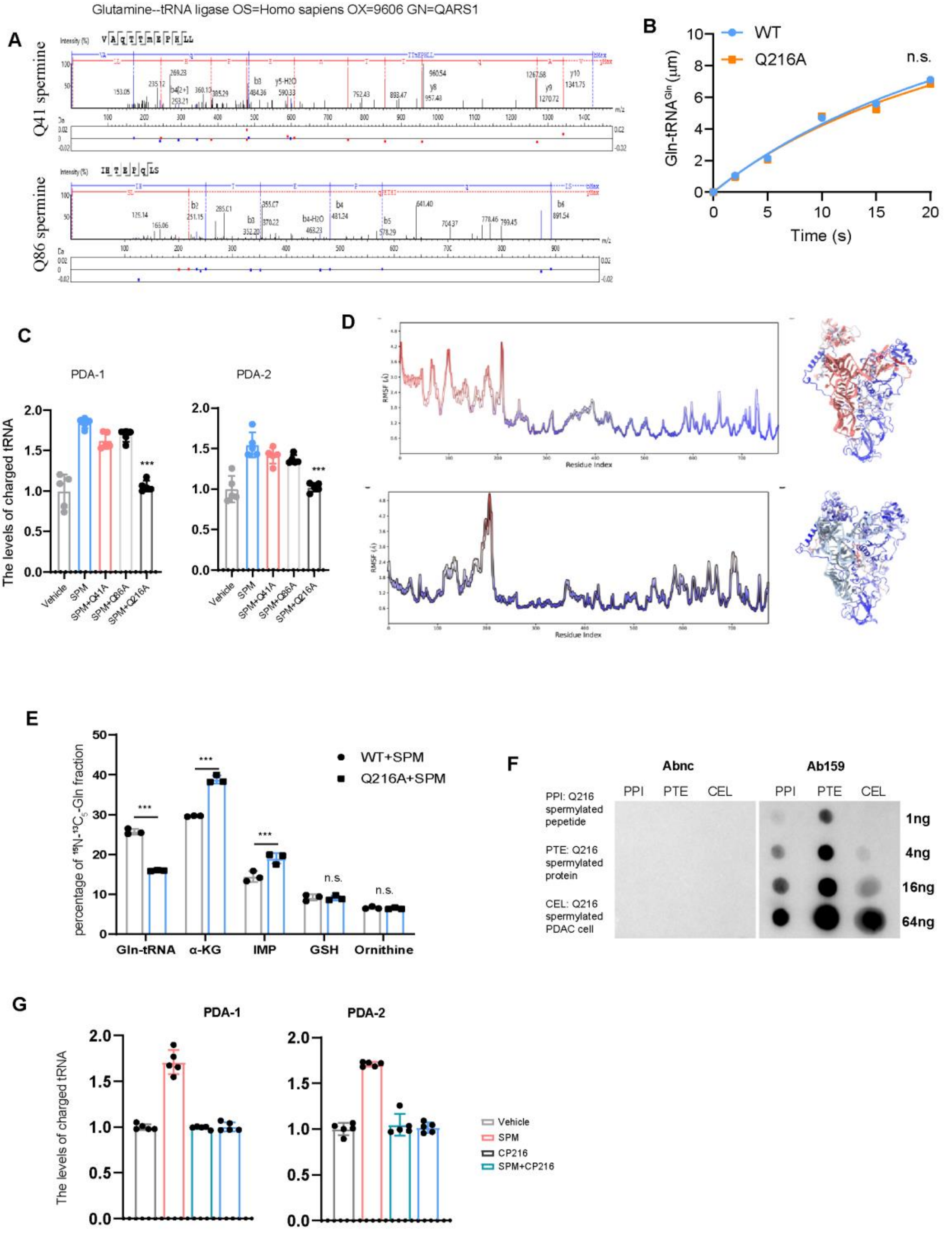


Supplement figure 3

**Q216 is the functionally dominant spermylation site controlling QARS activity and glutamine allocation. a,** Representative LC–MS/MS spectra supporting spermylation of QARS at Q41 and Q86; Q216 identification is shown in the main figure. **b,** Basal aminoacylation kinetics of recombinant QARS-WT and Q216A in the absence of spermylation, showing comparable Gln-tRNA formation. **c,** Relative abundance of charged Gln-tRNA in PDA-1 and PDA-2 cells expressing the indicated QARS site mutants following spermine treatment. **d,** Molecular-dynamics analysis of the QARS–tRNAGln complex, showing residue-level root-mean-square fluctuation profiles and representative structures for unmodified and Q216-spermylated QARS. **e,** U-13C5,15N2-glutamine allocation in spermine-treated QARS-WT and Q216A PDAC cells, showing label partitioning among Gln-tRNA, α-ketoglutarate (α-KG), IMP, GSH and ornithine. **f,** Dot-blot assessment of the specificity and sensitivity of the QARS Q216-spermylation-specific antibody (Ab159) compared with the control antibody (Abnc), using Q216-spermylated peptide, protein and cell-derived material across the indicated input amounts. **g,** Relative abundance of charged Gln-tRNA in PDA-1 and PDA-2 cells treated with vehicle, SPM, CP216, or SPM plus CP216. Data are mean ± s.d.; statistical analyses are described in Methods.

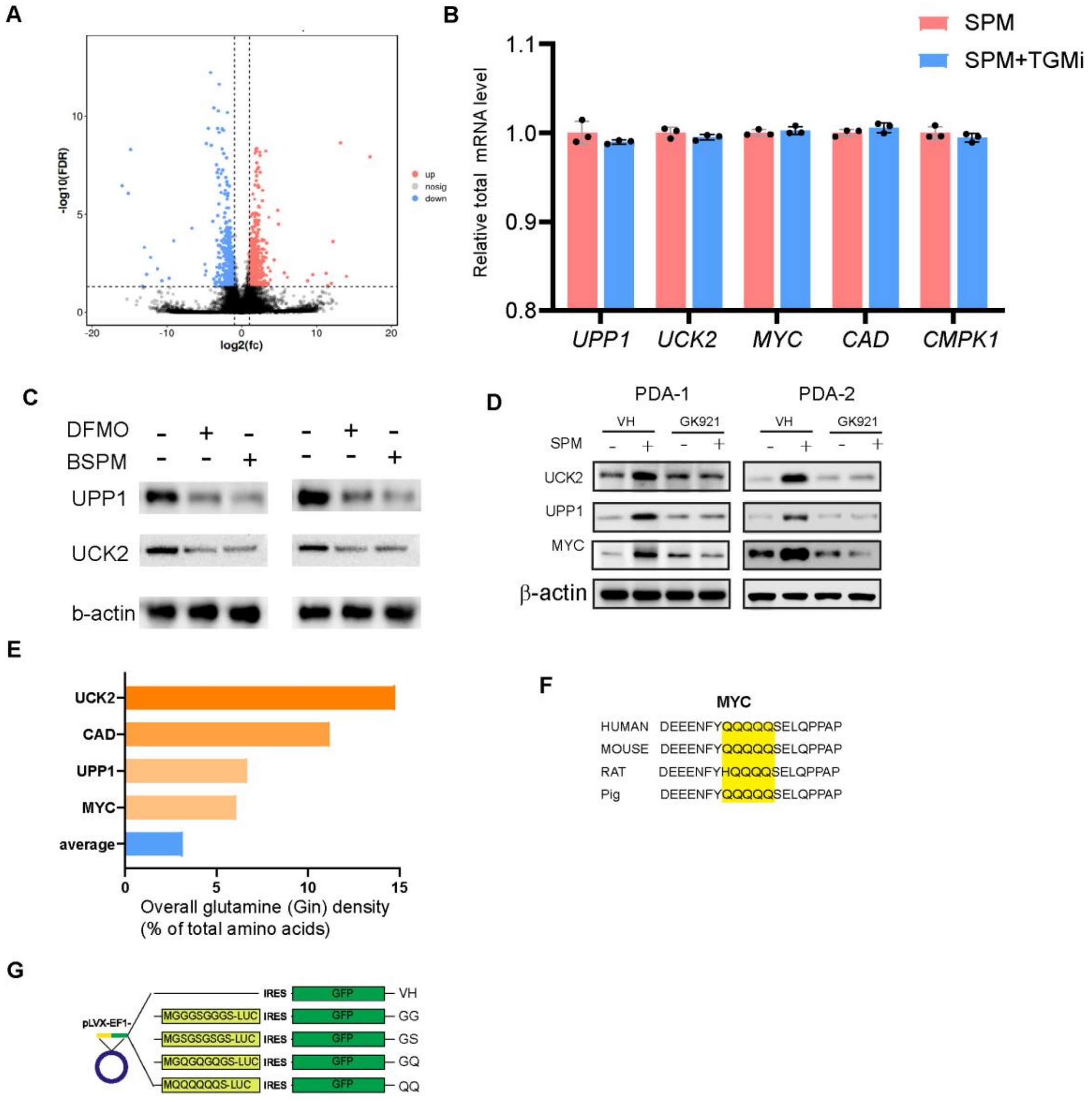


Supplement figure 4

**QARS spermylation selectively maintains translation of glutamine-codon-rich metabolic proteins. a,** Volcano plot of changes in translational efficiency following CP216-mediated blockade of QARS spermylation. **b,** Relative mRNA abundance of UPP1, UCK2, MYC, CAD and CMPK1 following SPM treatment with or without TGM2 inhibition, showing minimal changes at the transcript level. **c,** Immunoblot analysis of UPP1 and UCK2 in PDA-1 and PDA-2 cells following inhibition of spermine synthesis with DFMO or BSPM. **d,** Immunoblot analysis of UCK2, UPP1 and MYC in PDA-1 and PDA-2 cells treated with vehicle or SPM in the presence or absence of the TGM2 inhibitor GK921. **e,** Overall glutamine-residue density of UCK2, CAD, UPP1 and MYC compared with the proteome-wide average. **f,** Cross-species alignment of the glutamine-rich region of MYC; glutamine residues are highlighted. **g,** Schematic of the reporter constructs containing graded glutamine-codon content used to test codon-dependent translational responsiveness to QARS spermylation. Data are mean ± s.d.; statistical analyses are described in Methods.

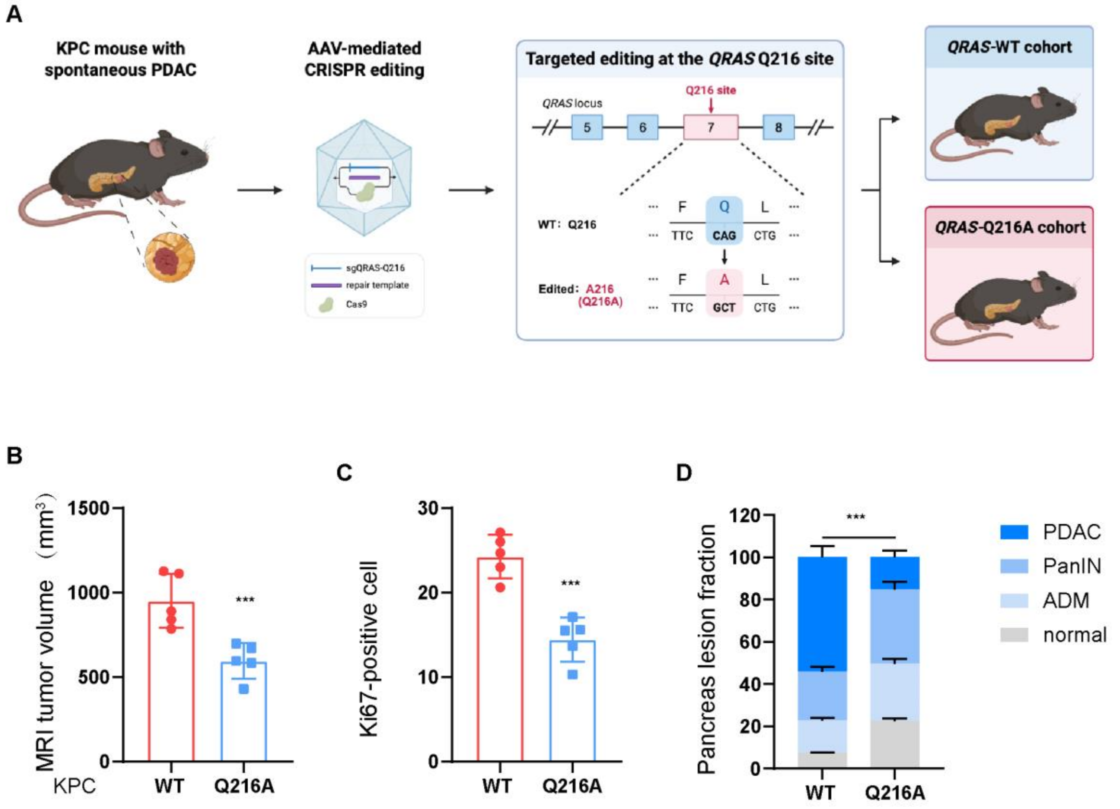


Supplement figure 5

**AAV-mediated QARS Q216A editing suppresses spontaneous PDAC progression in KPC mice.** **a,** Schematic of the AAV-mediated CRISPR strategy used to generate QARS-WT and QARS-Q216A KPC cohorts by targeted editing of the QARS Q216 locus. **b,** MRI-derived tumour-volume quantification in the AAV-edited QARS-WT and QARS-Q216A KPC cohorts. **c,** Quantification of Ki67-positive tumour cells in the corresponding KPC tumours. **d,** Histological composition of the pancreas, showing the relative fractions of normal tissue, acinar-to-ductal metaplasia (ADM), pancreatic intraepithelial neoplasia (PanIN) and PDAC. Data are mean ± s.d.; individual points represent biologically independent mice. Statistical analyses are described in Methods

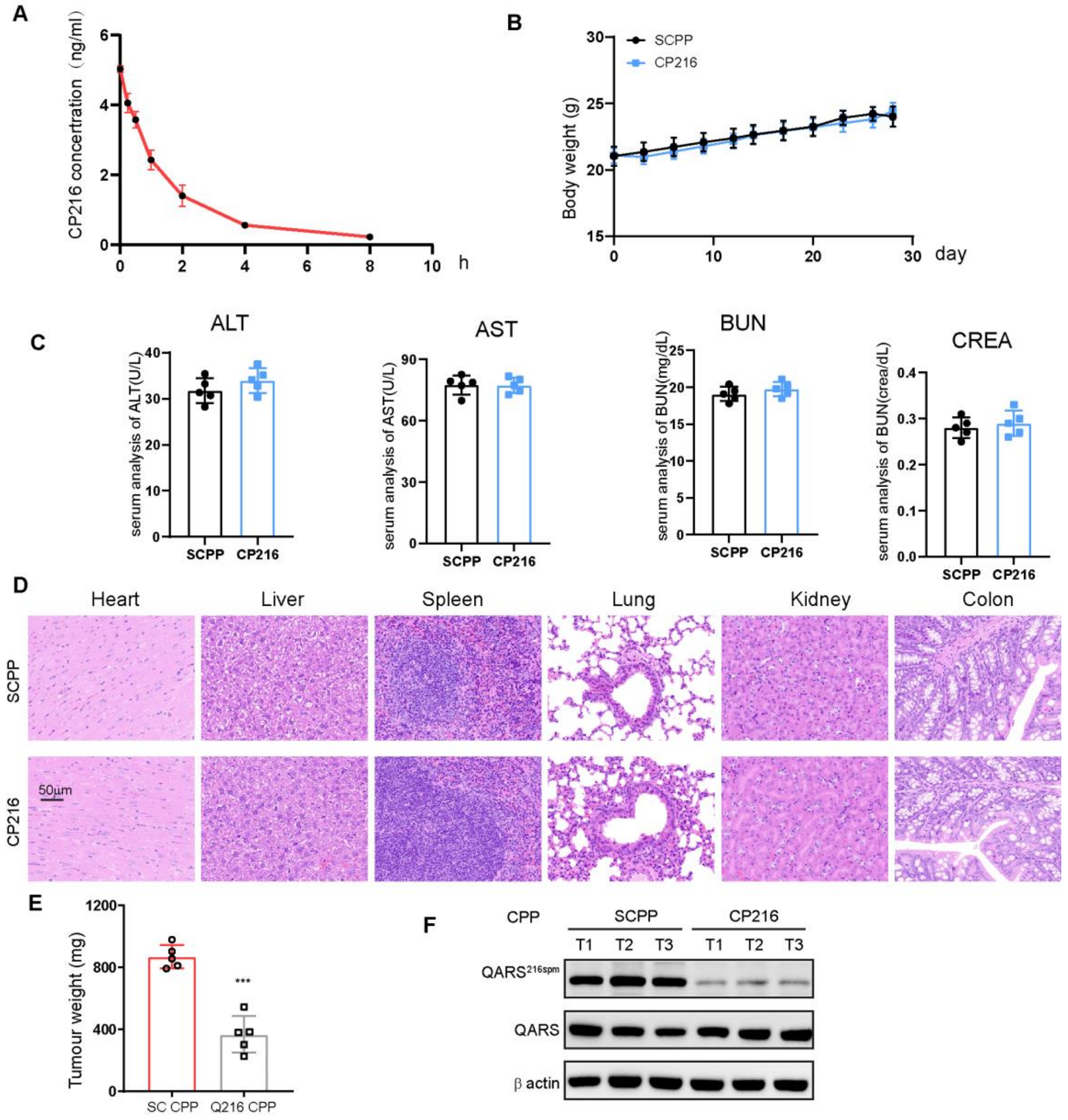


Supplement figure 6

**Pharmacokinetics, tolerability and in vivo target engagement of CP216. a,** Plasma concentration–time profile of CP216 after a single intraperitoneal dose of 10 mg kg−1 in six-week-old male C57BL/6J mice; n = 3 mice per time point. **b,** Body-weight trajectories during repeated treatment with scrambled control peptide (SC-CPP) or CP216 (10 mg kg−1, intraperitoneally, every 3 d for 4 weeks; n = 5 mice per group). **c,** Endpoint serum alanine aminotransferase (ALT), aspartate aminotransferase (AST), blood urea nitrogen (BUN) and creatinine (CREA) in the same treatment groups. **d,** Representative H&E staining of the indicated major organs after repeated SC-CPP or CP216 treatment. Scale bar, 50 μm. **e,** Endpoint tumour weight of PDA-1 subcutaneous xenografts treated with SC-CPP or CP216. **f,** Immunoblot analysis of QARS Q216 spermylation and total QARS in representative tumour samples following treatment with the indicated control peptide or CP216; β-actin is shown as a loading control. Data are mean ± s.d.; statistical analyses are described in Methods.

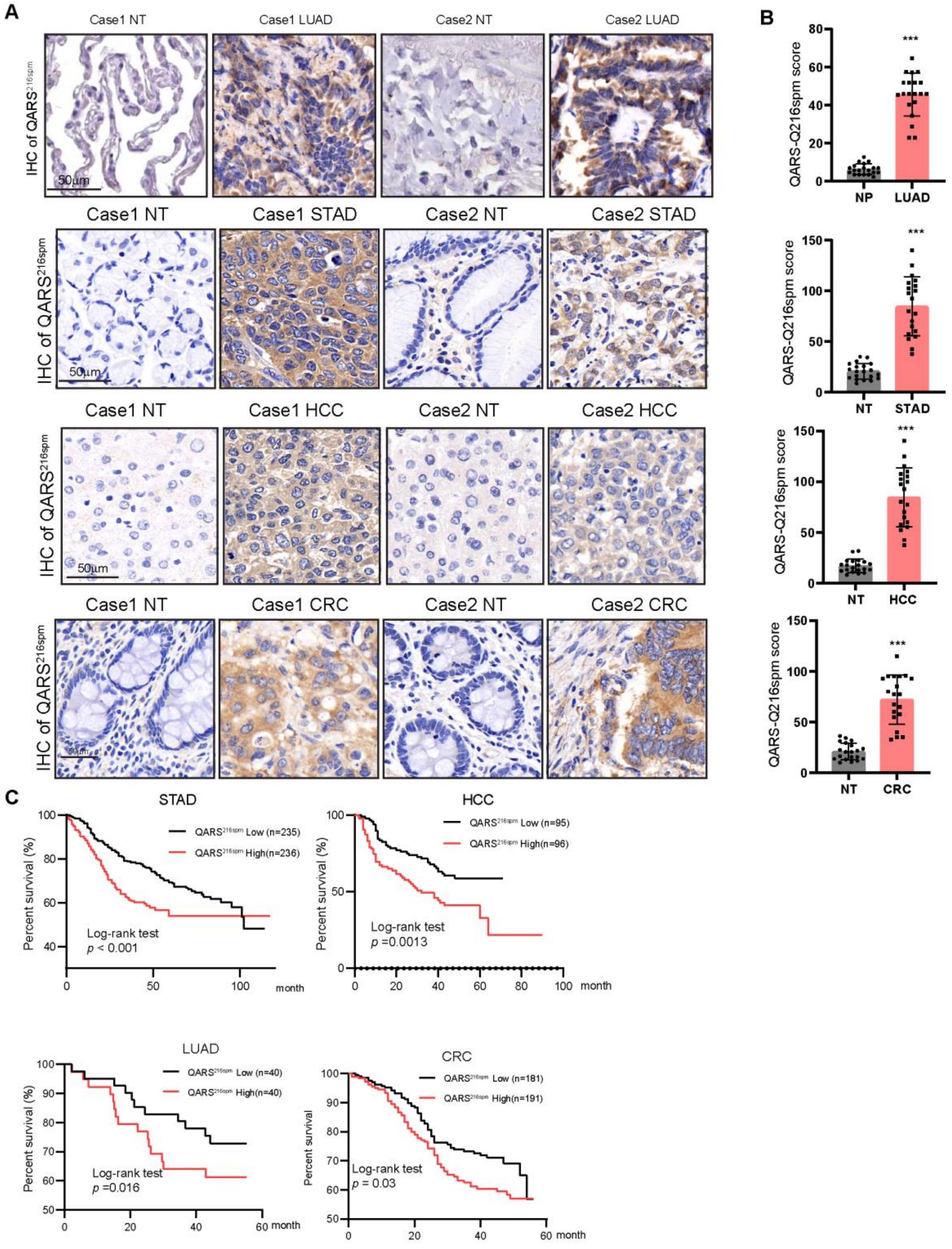


Supplement figure 7

QARS Q216 spermylation is elevated across multiple human malignancies and is associated with adverse outcome. a, Representative immunohistochemical staining of QARS Q216 spermylation in paired non-tumour and tumour tissues from lung adenocarcinoma (LUAD), stomach adenocarcinoma (STAD), hepatocellular carcinoma (HCC) and colorectal cancer (CRC). Scale bars,

50 μm. b, Quantification of QARS Q216 spermylation scores in the corresponding tumour and non-tumour cohorts. c, Kaplan–Meier overall-survival analyses stratified by low versus high QARS Q216 spermylation in STAD, HCC, LUAD and CRC. Group sizes are indicated in the corresponding plots; P values were calculated using two-sided log-rank tests. For quantitative IHC analyses, data are mean ± s.d.; statistical analyses are described in Methods.
